# Ethylene-mediated nitric oxide depletion pre-adapts plants to hypoxia stress

**DOI:** 10.1101/705194

**Authors:** Sjon Hartman, Zeguang Liu, Hans van Veen, Jorge Vicente, Emilie Reinen, Shanice Martopawiro, Hongtao Zhang, Nienke van Dongen, Femke Bosman, George W. Bassel, Eric J.W. Visser, Julia Bailey-Serres, Frederica L. Theodoulou, Kim H. Hebelstrup, Daniel J. Gibbs, Michael J. Holdsworth, Rashmi Sasidharan, Laurentius A.C.J. Voesenek

**Author notes:** Correspondence and requests for materials should be addressed to (L.A.C.J.V.), (R.S.) and (M.J.H.).

## Abstract

Timely perception of adverse environmental changes is critical for survival. Dynamic changes in gases are important cues for plants to sense environmental perturbations, such as submergence. In *Arabidopsis thaliana*, changes in oxygen and nitric oxide (NO) control the stability of ERFVII transcription factors. ERFVII proteolysis is regulated by the N-degron pathway and mediates adaptation to flooding-induced hypoxia. However, how plants detect and transduce early submergence signals remains elusive. Here we show that plants can rapidly detect submergence through passive ethylene entrapment and use this signal to pre-adapt to impending hypoxia. Ethylene can enhance ERFVII stability prior to hypoxia by increasing the NO-scavenger PHYTOGLOBIN1. This ethylene-mediated NO depletion and consequent ERFVII accumulation pre-adapts plants to survive subsequent hypoxia. Our results reveal the biological link between three gaseous signals for the regulation of flooding survival and identifies novel regulatory targets for early stress perception that could be pivotal for developing flood-tolerant crops.

## Introduction

The increasing frequency of floods due to climate change ^1^ has devastating effects on agricultural productivity worldwide ^2^. Due to restricted gas diffusion underwater, flooded plants experience cellular oxygen (O_2_) deprivation (hypoxia) and survival strongly depends on molecular responses that enhance hypoxia tolerance ^2,3^. In submerged plant tissues the limited gas diffusion causes passive ethylene accumulation. This rapid ethylene build-up can occur prior to the onset of severe hypoxia, making it a timely and reliable signal for submergence ^4,5^. In several plant species, ethylene regulates adaptive responses to flooding involving morphological and anatomical modifications that prevent hypoxia ^5^. Surprisingly, ethylene has so far not been linked to metabolic responses that reduce hypoxia damage. In addition, how plants detect and transduce early submergence signals to enhance survival remains elusive.

Here we show that plants can quickly detect submergence using passive ethylene accumulation and integrate this signal to acclimate to subsequent hypoxia. This ethylene-mediated hypoxia acclimation is dependent on enhanced ERFVII stability prior to hypoxia. We show that ethylene limits ERFVII proteolysis under normoxic conditions by increasing the NO-scavenger PHYTOGLOBIN1. Our results reveal a molecular mechanism that plants use to integrate early stress signals to pre-adapt to forthcoming severe stress.

## Results

### Early ethylene signalling enhances hypoxia acclimation

To unravel the spatial and temporal dynamics of ethylene signalling upon plant submergence, we monitored the nuclear accumulation of ETHYLENE INSENSITIVE 3 (EIN3) ^6–9^, an essential transcription factor for mediating ethylene responses. We show, through an increase in EIN3-GFP fluorescence signal, that ethylene is rapidly perceived (within 1-2 h) in *Arabidopsis thaliana* (hereafter Arabidopsis) root tips upon submergence (Supplementary Figure 1a-c). An ethylene or submergence pre-treatment of only 4 hours was sufficient to increase root meristem survival during subsequent hypoxia (<0.01% O_2_). These responses were abolished in ethylene signalling mutants or via chemical inhibition of ethylene action (Supplementary Figure 1d-e). Ethylene-induced adaptation to hypoxia was observed in both roots and shoots and was accompanied by a reduction in cellular damage in response to hypoxia (Fig. 1, Supplementary Figure 2 & 3). Furthermore, enhanced hypoxia tolerance after ethylene pre-treatment is conserved within Arabidopsis accessions and taxonomically diverse flowering plant species, although variation in capacity to benefit from an ethylene pre-treatment exists (Supplementary Figure 4; ^10^). These results demonstrate that ethylene enhances tolerance of multiple plant organs and species to hypoxia. Next, we aimed to unravel how early ethylene signalling leads to enhanced hypoxia tolerance in Arabidopsis root tips.

**Figure 1.**
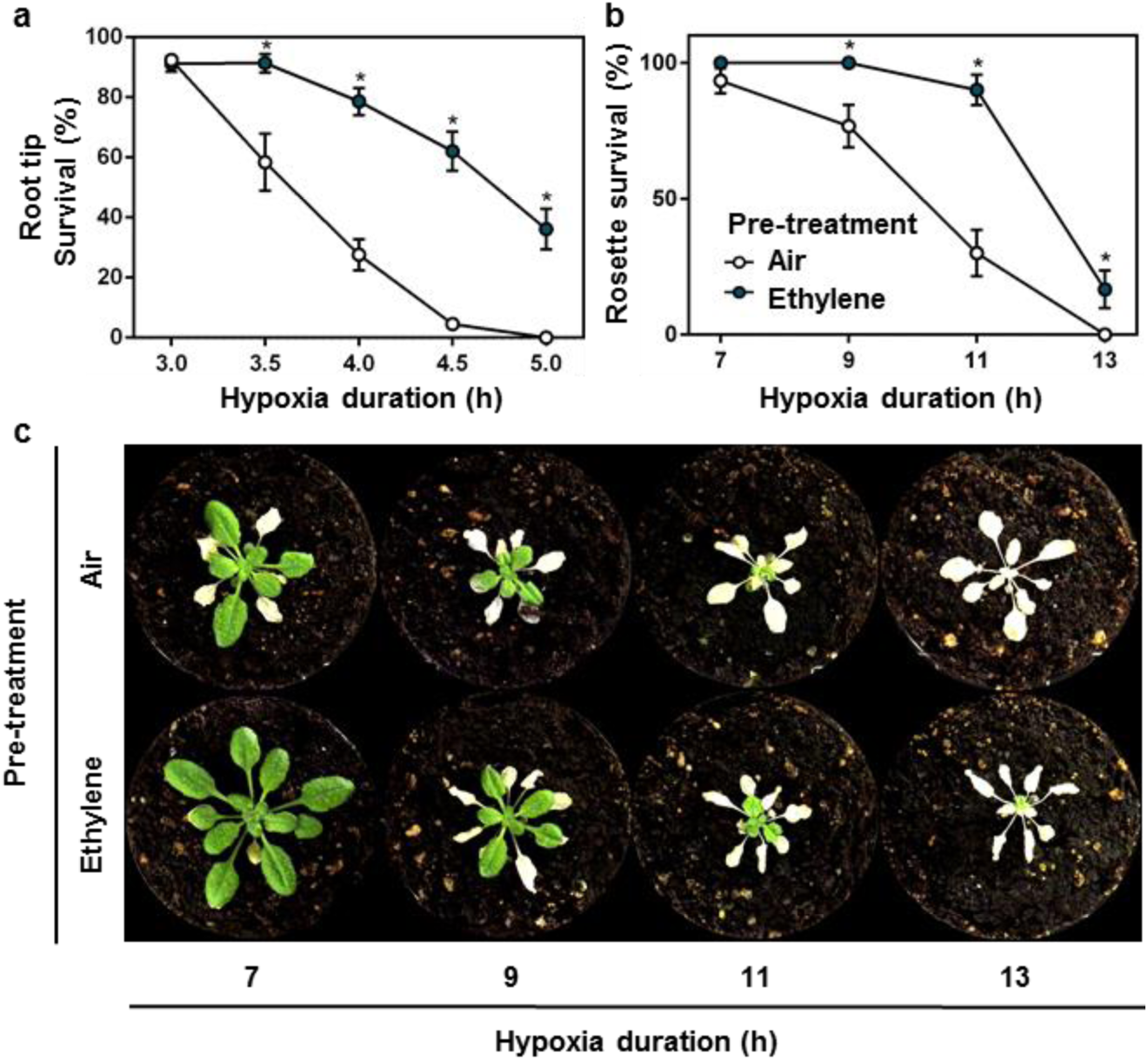
Ethylene pre-treatment enhances hypoxia tolerance. **(a, b)** Arabidopsis (Col-0) seedling root tip (a) and adult rosette (b) survival after 4 hours of air (white) or ~5μll^−1^ ethylene (blue) followed by hypoxia and recovery (3 days for root tips, 7 days for rosettes). Values are relative to control (normoxia) plants (mean ± sem). Asterisks indicate significant differences between air and ethylene (p<0.05, Generalized linear model, negative binomial error structure, n=4-8 lines consisting of ~23 seedlings (a), n=30 plants (b)). **(c)** Arabidopsis (Col-0) rosette phenotypes after 4 hours of pre-treatment (air/ ~5μll^−1^ ethylene) followed by hypoxia and 7 days recovery. All experiments were replicated at least 3 times.

### Ethylene stabilizes group VII Ethylene Response Factors

Hypoxia acclimation in plants involves the up-regulation of hypoxia adaptive genes that control energy maintenance and oxidative stress homeostasis ^11^. Interestingly, most of these genes were not induced by ethylene alone, but showed increased transcript abundance upon hypoxia following a pre-treatment with ethylene (Supplementary Figure 5). Hypoxia adaptive genes are regulated by group VII Ethylene Response Factor transcription factors (ERFVIIs) that are components of a mechanism that senses O_2_ and NO via the Cys-branch of the PROTEOLYSIS 6 (PRT6) N-degron pathway ^12–14^. ERFVIIs are degraded following oxidation of amino terminal (Nt-) Cysteine in the presence of oxygen and NO, catalysed by PLANT CYSTEINE OXIDASEs (PCOs) ^15^. The N-recognin E3 ligase PRT6 promotes degradation of oxidized ERFVIIs by the 26S proteasome ^16,17^. A decline in either O_2_ or NO stabilizes ERFVIIs, leading to transcriptional up-regulation of hypoxia adaptive genes and other environmental and developmental responses ^12,14,18,19^. The constitutively synthesized ERVIIs RELATED TO APETALA2.12 (RAP2.12), RAP2.2 and RAP2.3 redundantly act as the principal activators of many hypoxia adaptive genes ^20–22^. In contrast, HYPOXIA RESPONSIVE ERF1 (HRE1) and HRE2 function downstream of RAP-type ERFVIIs, being transcriptionally induced once hypoxia occurs ^23^. We investigated whether ethylene-induced hypoxia tolerance depends on the constitutively synthesized RAP-type ERFVIIs. Single loss-of-function mutants of *RAP2.12*, *RAP2.2* and *RAP2.3*, and the *hre1 hre2* double mutant, responded to ethylene pre-treatment similarly to their WT backgrounds (Supplementary Figure 6a). However, two independent *rap2.2 rap2.12* loss-of-function double mutants ^21^ showed no improved hypoxia tolerance after ethylene pre-treatment (Fig. 2a), while their WT background crosses did (Supplementary Figure 6b). In contrast, overexpression of a stable N-terminal variant of RAP2.12 ^22^, or inhibition of the PRT6 N-degron pathway in the *prt6-1* mutant ^12,24^ both enhanced hypoxia tolerance without an ethylene pre-treatment (Fig. 2a). These data indicate that ethylene-induced hypoxia tolerance occurs through the PRT6 N-degron pathway and redundantly involves at least RAP2.2 and RAP2.12.^21,22^.

**Figure 2.**
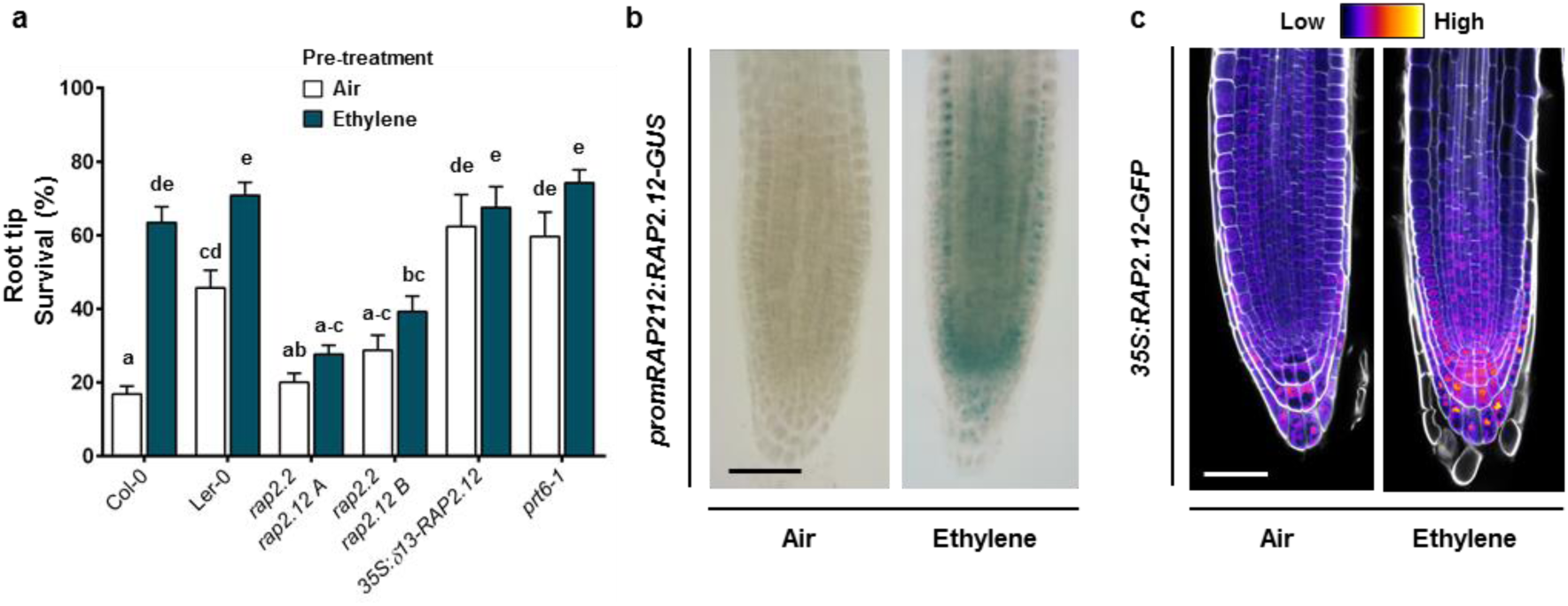
Ethylene-induced hypoxia tolerance is regulated by RAP-type ERFVIIs. **(a)** Seedling root tip survival of Col-0, Ler-0, *rap2.2 rap2.12* (2 independent lines in Col-0 x Ler-0 background), a constitutively expressed stable version of RAP2.12 and N-degron pathway mutant *prt6-1* after 4 hours air or ~5μll^−1^ ethylene followed by 4 hours of hypoxia and 3 days recovery. Values are relative to control (normoxia) plants (mean ± sem). Statistically similar groups are indicated using the same letter (p<0.05, 2-way ANOVA, Tukey’s HSD, n=20-28 lines consisting of ~23 seedlings). **(b)** and **(c)** Representative root tip images showing *promRAP2.12::RAP2.12-GUS* staining and confocal images of *35S::RAP2.12-GFP* intensity in root tips after 4 hours of air or ~5μll^−1^ ethylene. Cell walls were visualized using Calcofluor White stain (c). Scale bar of b and c is 50μm. All experiments were replicated at least 3 times.

We next explored how ethylene regulates *ERFVII* mRNA and protein abundance. Ethylene increased *RAP2.2*, *RAP2.3, HRE1* and *HRE2* transcripts in root tips and *RAP2.12*, *RAP2.2* and *RAP2.3* mRNAs in shoots (Supplementary Figure 6c-d). Visualization and quantification of RAP2.12 abundance using transgenic *promRAP2.12:RAP2.12-GUS* and *35S:RAP2.12-GFP* protein-fusion lines revealed that ethylene strongly increased RAP2.12 protein in meristematic zones of main and lateral root tips and shoots under normoxia (Fig. 2b-c, Supplementary Figure 6e-f). Since *35S::RAP2.12-GFP* is uncoupled from ethylene-triggered transcription, this suggests that ethylene limits ERFVII protein turnover. In root tips, this RAP2.12 stabilization appeared within nuclei across most cell types and was also independent of ethylene-enhanced RAP2.12 transcript abundance (Fig 2b-c, Supplementary Figure 6c, e-f). These data suggest that ethylene-enhanced ERFVII accumulation is regulated by post-translational processes.

### Ethylene limits ERFVII proteolysis through NO depletion

To investigate enhanced ERFVII stability under ambient O_2_, we studied the effect of ethylene on the expression of genes encoding PRT6 N-degron pathway enzymes or other mechanisms reported to influence ERFVII stability. In response to ethylene, none of these genes showed changes in transcript abundance (Supplementary Figure 7a-b). In addition, as both O_2_ and NO promote ERFVII proteolysis ^17^, and since ethylene was administered at ambient O_2_ conditions (21%; normoxia) and did not lead to hypoxia in desiccators (Supplementary Figure 7c), it is unlikely that hypoxia causes the observed ERFVII stabilization. Furthermore, while recent reports show that plants contain a hypoxic niche in shoot apical meristems and lateral root primordia ^25,26^, we did not observe enhanced hypoxia target gene expression in root tips exposed ethylene treatments (Supplementary Figure 5), ruling out ethylene-enhanced local hypoxia in these tissues.

Since NO was previously shown to control proteolysis of ERFVIIs and other Met_1_-Cys_2_ N-degron targets ^14,19,27^, we hypothesized that ethylene may regulate NO levels. Roots treated with the NO probe 4-Amino-5-Methylamino-2’,7’-Difluorofluorescein (DAF-FM) Diacetate ^28^ revealed an ethylene-induced depletion in fluorescence, indicating that ethylene mediates NO levels (Fig. 3a-b). Next, we investigated whether this decline in NO was required for RAP-type ERFVII stabilization. Both ethylene and the NO-scavenging compound 2-(4-Carboxyphenyl)-4,4,5,5-Tetramethylimidazoline-1-oxyl-3-oxide (cPTIO) led to increased RAP2.12 and RAP2.3 stability under normoxia (Fig. 3c-e). However, the ethylene-mediated increase in RAP2.12 and RAP2.3 stability was abolished when an NO pulse was applied concomitantly confirming a role for NO depletion in ethylene-triggered ERFVII stabilization of both these RAPs during normoxia. Application of hypoxia after pre-treatments resulted in stabilization of RAP2.12 and RAP2.3, demonstrating that the plants were viable and the PRT6 N-degron pathway could still be impaired (Fig. 3c-e, Supplementary Figure 7d). These data together illustrate that both RAP2.12 and RAP2.3 depend on ethylene-mediated NO-depletion to promote their stability. The functional consequences of ethylene-induced NO-dependent RAP2.12 stabilization for hypoxia acclimation were studied in a root meristem survival assay. Ethylene pre-treatment enhanced hypoxia survival, which was largely abolished by an NO pulse (Fig. 3f). Furthermore, pre-treatment with cPTIO to scavenge intracellular NO before hypoxia resulted in increased survival in the absence of ethylene. In genotypes lacking RAP2.12 and RAP2.2 or overexpressing a stable N-terminal variant of RAP2.12, neither ethylene nor NO manipulation had any effect on subsequent hypoxia survival (Fig. 3f). These results demonstrate that local NO removal, via cPTIO or as a result of elevated ethylene, is both essential and sufficient to enhance RAP2.12 and RAP2.3 stability during normoxia, and that increased hypoxia tolerance conferred by ethylene strongly depends on NO-mediated stabilization of RAP2.12 and RAP2.2 prior to hypoxia.

**Figure 3.**
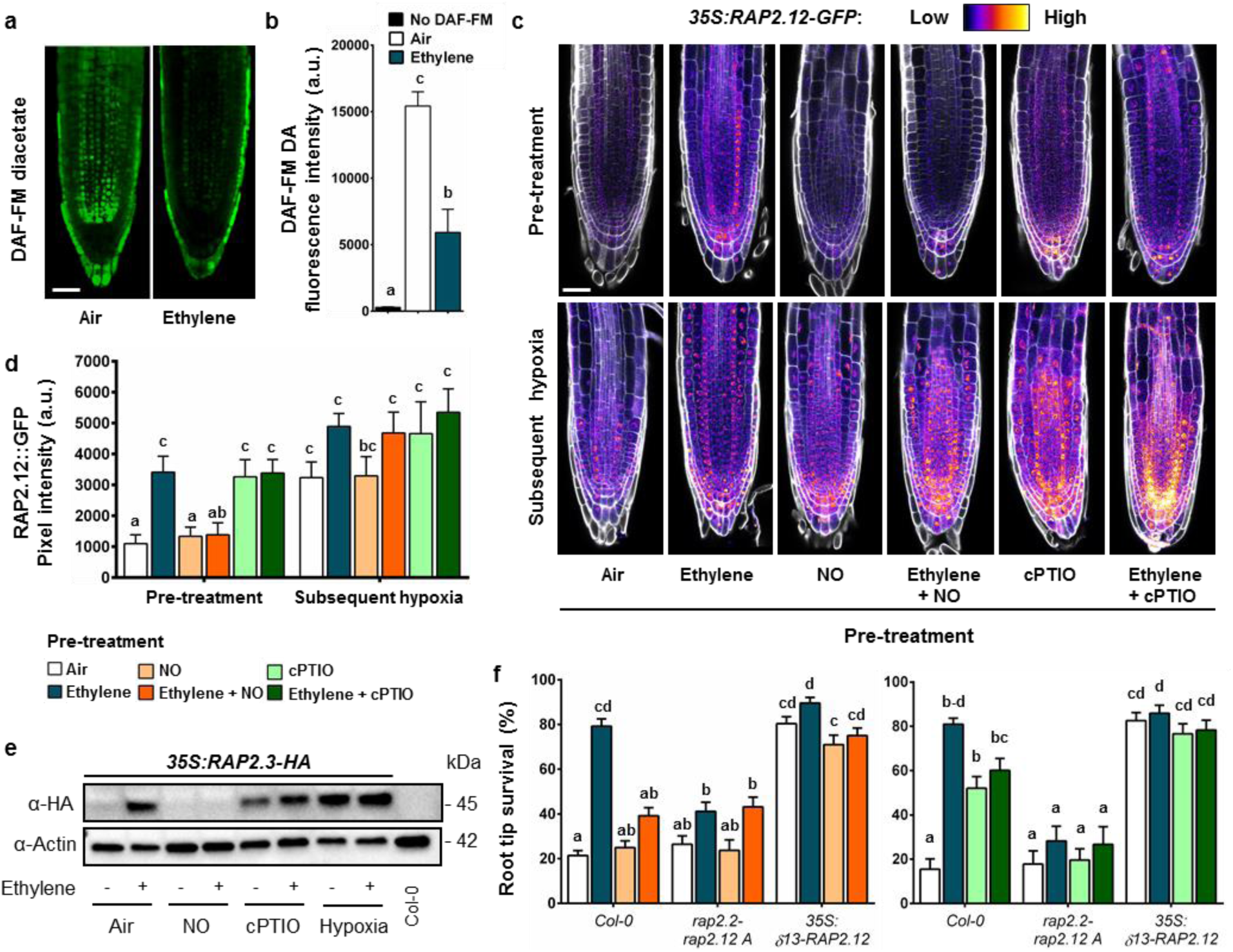
Ethylene impairs NO levels leading to ERFVII stability and enhanced hypoxia survival. **(a, b)** Representative confocal images visualizing **(a)** and quantifying **(b)** NO, using fluorescent probe DAF-FM diacetate in Col-0 seedling root tips after 4 hours of air or ~5μll^−1^ ethylene (scale bar = 50μm). (Letters indicate significant differences (1-way ANOVA, Tukey’s HSD, n=5). **(c, d)** Representative confocal images visualizing (c) and quantifying (d) *35S::RAP2.12-GFP* intensity in seedling root tips after indicated pre-treatments and subsequent hypoxia (4h). Cell walls were visualized using Calcofluor White stain (scale bar = 50μm). (Letters indicate significant differences (p<0.05, 2-way ANOVA, Tukey’s HSD, n=5-7). **(e)** RAP2.3 protein levels in 35S::MC-RAP2.3-HA seedlings (Col-0 background) after indicated treatments. **(f)** Seedling root tip survival of Col-0, *rap2.2 rap2.12* line *A* mutants and an over-expressed stable version of RAP2.12 after indicated pre-treatments followed by hypoxia (4h) and 3 days recovery. Values are relative to control (normoxia) plants. Letters indicate significant differences (p<0.05, 2-way ANOVA, Tukey’s HSD, n=12 rows consisting of ~23 seedlings). All data shown are mean ± sem. All experiments were replicated at least 3 times, except for c, d and f (2 times).

### Ethylene-mediated NO depletion is regulated by PHYTOGLOBIN1

The question remained how ethylene regulates NO levels under normoxia. NO metabolism in Arabidopsis is mainly regulated by NO biosynthesis via NITRATE REDUCTASE (NR)-dependent nitrite reduction and NO-scavenging by three non-symbiotic phytoglobins (PGBs) ^29–31^. Ethylene led to small increases in *NR1* and *NR2* mRNA levels, but this did not influence total NR activity (Supplementary Figure 8a, b&e). In contrast, transcript abundance of *PGB1*, the most potent NO-scavenger ^31^, increased rapidly in root tips and shoots after ethylene treatment (Supplementary Figure 8a-c). Importantly, *PGB1* (a hypoxia-adaptive gene regulated by ERFVIIs) was still up-regulated by ethylene during normoxia in *rap2.2 rap2.12* mutant lines (Supplementary Figure 8d). To study the effect of ethylene-induced *PGB1* levels on NO metabolism, ERFVII stabilization, hypoxia-adaptive gene expression and hypoxia tolerance, we identified a T-DNA insertion line (*SK_058388*; hereafter *pgb1-1*). In *pgb1-1* the T-DNA is located 300 bp upstream of the *PGB1* start codon (Supplementary Figure 9a-b). In wild-type plants, both ethylene and hypoxia treatment enhanced *PGB1* transcript and protein accumulation (Fig. 4a-b). In *pgb1-1*, *PGB1* transcript levels were reduced, and ethylene did not increase *PGB1* transcript or protein abundance, whereas hypoxia only affected transcript abundance slightly (Fig. 4a-b). A faint band of lower molecular weight than expected for PGB1 (18 kDa) was observed in some *pgb1-1* samples, but did show any clear treatment effect (Fig 4b). Together these data illustrate that the T-DNA insertion in the promoter of *pgb1-1* uncouples *PGB1* expression from ethylene regulation. Conversely, a *35S:PGB1* line had constitutively elevated *PGB1* transcript and protein levels (Fig 4a-b, ^31^). Importantly, both *pgb1-1* and *35S:PGB1* showed mostly similar ethylene responses in abundance of perception (*ETR2*) and biosynthesis (*ACO1*) transcripts compared to wild-type during normoxia (Supplementary Figure 10), indicating that ethylene biosynthesis and signalling are unlikely to be affected.

**Figure 4.**
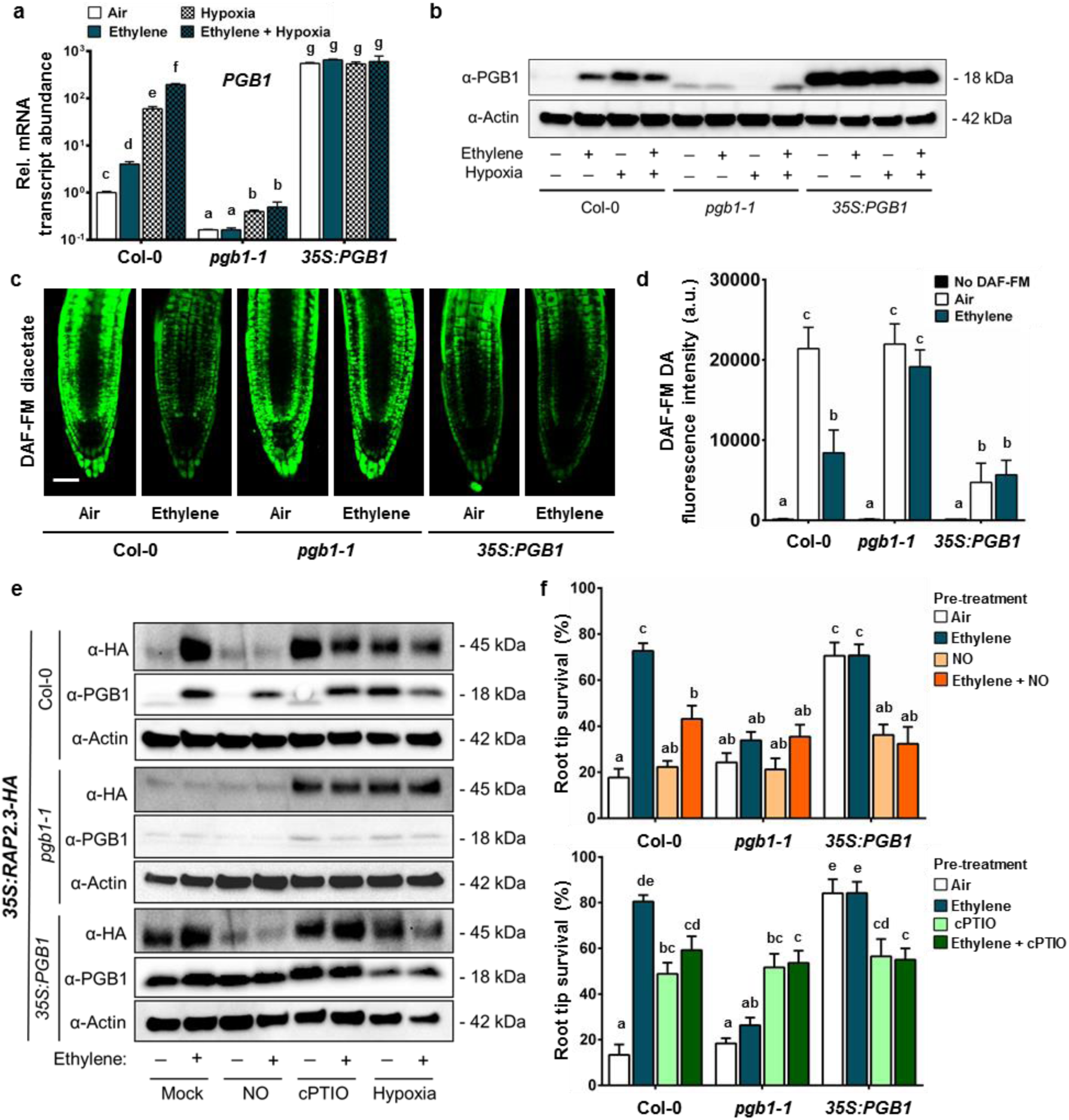
Ethylene mediates NO levels, ERFVII stability and hypoxia survival through PHYTOGLOBIN1. **(a)** Relative transcript abundance of *PGB1* in root tips of Col-0*, pgb1-1* and *35S::PGB1* after 4 h air or ~5μll^−1^ ethylene followed by (4h) hypoxia. Values are relative to Col-0 air treated samples. Letters indicate significant differences (p<0.05, 2-way ANOVA, n=3 replicates of ~200 root tips each). **(b)** PGB1 protein levels in Col-0 *, pgb1-1* and *35S::PGB1* root tips after 4 h air or ~5μll^−1^ ethylene followed by (4h) hypoxia. **(c, d)** Representative confocal images visualizing (C) and quantifying (D) NO using fluorescent probe DAF-FM diacetate in Col-0, *pgb1-1* and *35S:PGB1* seedling root tips after 4h air or ~5μll^−1^ ethylene (scale bar = 50μm). Letters indicate significant differences (p<0.05, 2-way ANOVA, Tukey’s HSD, n=5) **(e)** RAP2.3 and PGB1 protein levels in *35S::MC-RAP2.3-HA* (in Col-0, *pgb1-1* and *35S::PGB1* background) seedling root tips after indicated pre-treatments and subsequent hypoxia (4h). **(f)** Seedling root tip survival of Col-0*, pgb1-1* and *35S::PGB1* after indicated pre-treatments followed by 4 h hypoxia and 3 days recovery. Values are relative to control (normoxia) plants. Letters indicate significant differences (p<0.05, 2-way ANOVA, Tukey’s HSD, n=12 rows of ~23 seedlings). All data shown are mean ± sem. All experiments were replicated at least 2 times.

The ethylene-mediated NO decline observed in wild-type root tips was fully abolished in *pgb1-1*, demonstrating the requirement of *PGB1* induction for local NO removal upon ethylene exposure (Fig. 4c-d). Moreover, lack of NO removal by ethylene in *pgb1* resulted in the inability to stabilize RAP2.3 levels and reduced hypoxia survival (Fig. 4e-f). These effects could be rescued by restoration of NO-scavenging capacity using cPTIO (Fig. 4f). In addition, the reduced ethylene-induced hypoxia tolerance in *pgb1-1* was also accompanied by an absence of enhanced hypoxia adaptive gene expression after an ethylene pre-treatment (Supplementary Figure 10). In contrast, *35S:PGB1* showed constitutively low NO levels in root tips (Fig. 4c, d, ^31^), and increased RAP2.3 stability under normoxia (Fig. 4e). Moreover, ectopic *PGB1* over-expression enhanced hypoxia tolerance without an ethylene pre-treatment, but this effect was abolished by an NO pulse (Fig. 4f). Elevated mRNA levels for several hypoxia adaptive genes accompanied this constitutive hypoxia tolerance in *35S:PGB1* root tips (Supplementary Figure 10). These results demonstrate that active reduction of NO levels by ethylene-induced *PGB1* prior to hypoxia can precociously enhance ERFVII stability to prepare cells for impending hypoxia.

## Discussion

We show that plants have the remarkable ability to detect submergence quickly by passive ethylene entrapment and use this signal to acclimate to forthcoming hypoxic conditions. The early ethylene signal prevents N-degron targeted ERFVII proteolysis by increased production of the NO-scavenger PGB1 and in turn primes the plant’s hypoxia. Interestingly, while ethylene signalling prior to hypoxia leads to nuclear stabilization of RAP2.12 in root meristems (Fig. 2b-c, 3c), it does not trigger accumulation of most hypoxia adaptive gene transcripts until hypoxia occurs (Supplementary Figure 5). Apparently, stabilization of ERFVIIs alone is insufficient to trigger full activation of hypoxia-regulated gene transcription and additional hypoxia-specific signals such as altered ATP and/or Ca^2+^ levels are required ^32–34^. The possible existence of undiscovered plant O_2_ sensors was recently discussed and could potentially fulfil this role ^35^. Furthermore, the current discovery of ethylene-mediated stability of ERFVIIs paves the way towards unravelling how ethylene could influence the function of the other recently discovered PRT6 N-degron pathway targets VERNALIZATION2 (VRN2) and LITTLE ZIPPER2 (ZPR2)^25,27^.

This study shows that PGB1 is a key intermediate, linking ethylene signalling, via regulated NO removal, to O_2_ sensing and hypoxia tolerance. This mechanism also provides a molecular explanation for the protective role of PGB1 during hypoxia and submergence described in prior studies ^31,36–38^. Natural variation for ethylene-induced hypoxia adaptation was also observed in wild species and correlated with *PGB1* induction ^10^. Our discovery provides an explanation for this natural variation and could be instrumental in enhancing conditional flooding tolerance in crops via manipulation of ethylene responsiveness of *PGB1* genes. In these modified plants, rapid passive ethylene entrapment upon flooding would increase PGB1 levels and pre-adapt crops to later occurring hypoxia stress.

## Methods

### Plant material and growth conditions

#### Plant material

*Arabidopsis thaliana* seeds of ecotypes Col-0, Cvi-0, C24 and mutants *ein2-5* and *ein3eil1-1* ^39,40^ were obtained from the Nottingham Arabidopsis Stock Centre. Seeds of *pgb1-1* (SALK_058388) were obtained from the Arabidopsis Biological Resource Center and the molecular characterization of this line is described in Fig. 4a-b and Supplementary Figure 9. Other germlines used in this study were kindly provided by the following individuals: Ler-0, *rap2.2-5* (Ler-0 background, AY201781/GT5336), *rap2.12-2* (SAIL_1215_H10), *rap2.2-5rap2.12-A* and *-B* (mixed Ler-0 and Col-0 background) from Prof. Angelika Mustroph ^21^, University Bayreuth, Germany; *35S:δ13-RAP2.12-GFP* and *35S:RAP2.12-GFP* from Prof. Francesco Licausi, University of Pisa, Italy ^13^; and *35S:EIN3-GFP* (*ein3eil1* mutant background) from Prof. Shi Xiao, Sun Yat-sen University, China ^7^. The *35S:PGB1*, *35S::RAP2.3-HA* transgenic lines, as well as *prt6-1* (SAIL_1278_H11), *rap2.3-1* (SAIL_1031_D10) and *hre1-1hre2-1* (SALK_039484 + SALK_052858) mutants were described previously ^12,14,41^. Barley seeds were obtained from Flakkebjerg Research Center Seed Stock (Aarhus University). Additional mutant combinations used in this study were generated by crossing, and all lines were confirmed by either conventional genotyping PCRs and/or antibiotic resistance selection (Primer and additional info in Table S1).

#### Growth conditions adult rosettes

Arabidopsis seeds were placed on potting soil (Primasta) in medium sized pots and stratified at 4°C in the dark for at least 3 days. Pots were then transferred to a growth chamber for germination under short day conditions (8:00 – 17:00, T= 20°C, Photon Flux Density = ~150 μmol m^−2^s^−1^, RH= 70%). After 7 days, seedlings were transplanted individually into single pots (70ml) that were filled with the similar potting soil (Primasta). Plants continued growing under identical short day conditions and were watered automatically to field capacity. Per genotype, homogeneous groups of 10-leaf-stage plants were selected and randomized over treatment groups for phenotypic and molecular analysis under various treatments. Plants used for these experiments were transferred back to the same conditions after treatments to recover for 7 days.

#### Growth conditions seedlings

Seeds were vapor sterilized by incubation with a beaker containing a mixture of 50 ml bleach and 3 ml of fuming HCl in a gas tight desiccator jar for 3 to 4 hours. Seeds were then individually transplanted in (2 or 3) rows of 23 seeds on sterile square petri dishes containing 25 ml autoclaved and solidified ¼ MS, 1% plant agar without additional sucrose. Petri dishes were sealed with gas-permeable tape (Leukopor, Duchefa) and stratified at 4°C in the dark for 3 to 4 days. Seedlings were grown vertically on the agar plates under short day conditions (9:00 – 17:00, T= 20°C, Photon Flux Density = ~120 μmol m^−2^s^−1^, RH= 70%) for 5 days for *Arabidopsis thaliana*, and 7 days for *Solanum lycopersicum* (Tomato, Moneymaker), *Solanum dulcamara* and *Arabidopsis lyrata* before phenotypic and/or molecular analysis under various treatments. For *Hordeum vulgare* (Barley, both ssp. Golden Promise and landrace Heimdal) seedlings were grown on agar in sterile tubs and were 3 days old before phenotypic analysis.

### Construction of transgenic plants

The *promRAP2.12:MC-RAP2.12-GUS* protein fusion lines were constructed by amplifying the genomic sequence capturing 2 kb of sequence upstream of the translational start site, and removing the stop codon using the following primers: RAP2.12-fwd GGGGACAAGTTTGTACAAAAAAGCAGGCTATTCAGATTGGATCGTGACATG and RAP2.12-rev GGGGACCACT TTGTACAAGAAAGCTGGGTAGAAGACTCCTCCAATCATGGAAT. The PCR product was GATEWAY cloned into pDNR221 through a BP reaction, then transferred to pGWB433 creating an in-frame C-terminal fusion to the GUS reporter protein ^42^.

### Experimental setup and (pre-)treatments

#### Ethylene treatments

Lids of the agar plates of the vertically grown seedlings were removed during all (pre-) treatments and plates were placed vertically into glass desiccators (22.5 L volume). Air (control) and ~5μll^−1^ ethylene (pre-) treatments (by injection with a syringe) were applied at the start of the light period (9:00 for seedlings, 8:00 for adult rosettes) and were performed by keeping the seedlings/plants in air-tight closed glass desiccators under low light conditions (T= 20°C, Light intensity= ~3-5 μmol m^−2^s^−1^) for 4 hours. Ethylene concentrations in all desiccators were verified by gas chromatography (Syntech Spectras GC955) at the beginning and end of the pre-treatment.

#### Hypoxia treatments

After 4 hours of any pre-treatment plants/seedlings were flushed with oxygen-depleted air (humidified 100% N_2_ gas) at a rate of 2.00 l/min under dark conditions to limit oxygen production by photosynthesis. Oxygen levels generally reached 0.00% oxygen within 40 minutes of the hypoxia treatment as measured with a Neofox oxygen probe (Ocean optics, Florida, USA) (Supplementary Figure 7c). Control plants and seedlings were flushed with humidified air condition for the duration of the hypoxia treatment in the dark. Hypoxia treatment durations varied depending on the developmental stage and plant species and are specified in the appropriate figure legends.

#### Nitric oxide

Just before application, pure NO gas was diluted in small glass vials with pure N_2_ gas to minimize the oxidation of NO gas. Diluted NO gas was injected with a syringe into the air and ethylene treated desiccators at a final concentration of 10 ull^−1^ NO, 1 hour prior to the end of the (pre-)treatment.

#### c-PTIO

Treatments with the NO-scavenger 2-(4-Carboxyphenyl)-4,5-dihydro-4,4,5,5-tetramethyl-1H-imidazol-1-yloxy-3-oxide potassium salt (c-PTIO salt, Sigma Aldrich, Darmstadt, Germany) were performed 1 hour prior to ethylene treatments to allow for treatment combinations. Droplets of 5μl c-PTIO solution (250μM in autoclaved liquid ¼ MS) or mock solution (autoclaved liquid ¼ MS) were pipetted onto each individual root tip.

#### 1-MCP

Seedlings were placed in closed glass desiccators (22.5l volume) and gassed with 5μll^−1^ 1-MCP (Rohmand Haas) for 1 hour prior to other (pre-) treatments.

#### Submergence

For submergence (pre-) treatments, the plates of vertically grown seedlings were placed horizontally and were carefully filled with autoclaved tap water until the seedlings were fully submerged.

### Hypoxia tolerance assays

#### Adult rosette plants

10-leaf stage plants received ethylene and air pre-treatments followed by several durations of hypoxia and were subsequently placed back under short day growth chamber conditions to recover. After 7 days of recovery survival rates and biomass (fresh and dry weight of surviving plants) were determined.

#### Root tip survival of seedlings

5-day old seedlings grown vertically on agar plates received pre-treatments (described above) followed by several durations of hypoxia (generally 4 hours for mutant analysis). After the hypoxia treatment, agar plates were closed and sealed again with Leukopor tape and the location of root tips was marked at the back of the agar plate using a marker pen (0.8mm fine tip). Plates were then placed back vertically under short day growth conditions for recovery. After 3-4 days of recovery, seedling root tips were scored as either alive or dead based on clear root tip re-growth beyond the line on the back of the agar plate. Primary root tip survival was calculated as the percentage of seedlings that showed root tip re-growth out of a row of (maximally) 23 seedlings. Root tip survival was expressed as relative survival compared to control plates that received similar pre-treatments but no hypoxia. For *Solanum lycopersicum* (Tomato, Moneymaker), *Solanum dulcarama* and *Arabidopsis lyrata* methods were similar as described above, but seedlings were 7 days old. For *Hordeum vulgare* (Barley, both ssp. Golden Promise and landrace Heimdal) seedlings were only 3 days old and received 20 hours of hypoxia before scoring survival of whole seedlings after 3 days of recovery.

#### Evans blue staining for cell viability in root tips

Arabidopsis seedlings were taken for root cell integrity analysis by Evans blue staining after air and ethylene pre-treatments followed by both hypoxia and post-hypoxia time-points. Seedlings were incubated in 0.25% aqueous Evans blue staining solution for 15 minutes in the dark, subsequently washed three times with Milli-Q water to remove excess dye and finally imaged using light microscopy (OLYMPUS BX50WI, 10x objective). Evans blue area and pixel intensity of the microscopy images was analyzed using ICY software (http://icy.bioimageanalysis.org/), by quantifying the mean pixel intensity of the red (ch0) and blue (ch2) channels of the tissues of interest, and expressed as Blue/Red pixel intensity.

### RNA extraction and quantification, cDNA synthesis and RT-qPCR

Adult rosette (2 whole rosettes per sample), whole seedling (~20 whole seedlings) or seedling root tip (~500 root tips) samples were harvested by snap freezing in liquid nitrogen. Total sample RNA was extracted from frozen pulverized tissue using the RNeasy Plant Mini Kit protocol (Qiagen, Dusseldorf, Germany) with on-column DNAse treatment Kit (Qiagen, Dusseldorf, Germany) and quantified using a NanoDrop ND-1000 UV-Vis spectrophotometer (Nanodrop Technology). Single-stranded cDNA was synthesized from 500 ng RNA using random hexamer primers (Invitrogen, Waltham, USA). qRT-PCR was performed using the Applied Biosystems ViiA 7 Real-Time PCR System (Thermo Fisher Scientific) with a 5μl reaction mixture containing 2.5μl 2× SYBR Green MasterMix (Bio-Rad, Hercules, USA), 0.25μL of both 10μM forward and reverse primers and 2μl cDNA (5ng/μl). Average sample CT values were derived from 2 technical replicates. Relative transcript abundance was calculated using the comparative CT method ^43^, fold change was generally expressed as fold change relative to air treated samples of Col-0. *ADENINE PHOSPHORIBOSYL TRANSFERASE 1* (*APT1*) was amplified, stable in all treatments and used as a reference gene. Primers used for RT-qPCR can be found in Table S2.

### Histochemical staining for GUS activity

Seedlings of *promRAP2.12:RAP2.12-GUS* (10 days old) were harvested in GUS solution (100mM NaPO4 buffer, pH 7.0, 10mM EDTA, 2mM X-Gluc, 500 μM K3Fe(CN)6 and 500 μM K4Fe(CN)6) directly after (indicated in figure legend) treatments, vacuum infiltrated for 15 minutes and incubated for 2 days at 37°C before de-staining with 70% ethanol. Seedlings were kept and mounted in 50% glycerol and analyzed using a Zeiss Axioskop2 DIC (differential interference contrast) microscope (10× DIC objective) or regular light microscope with a Lumenera Infinity 1 camera. GUS pixel intensity of the microscopy images was analyzed using ICY software (http://icy.bioimageanalysis.org/), by quantifying the pixel intensity of the red (ch0) and blue (ch2) channels of the tissues of interest relative to the respective channel background values of these images. GUS intensity of all treatments was expressed relatively to the Air-treated controls.

### Protein extraction, SDS-PAGE and Western Blotting

Protein was extracted on ice for 30 minutes from pulverized snap frozen samples in modified RIPA lysis buffer containing 50 mM HEPES-KOH pH (7.8), 100 mM KCl, 5 mM EDTA (pH 8), 5 mM EGTA (pH 8), 50 mM NaF, 10% (v/v) glycerol, 1% (v/v) IGEPAL, 0.5% (w/v) deoxycholate, 0.1% (w/v) SDS, 1 mM Na3VO4 (sodium orthovanadate), 1 mM PMSF, 1x proteinase inhibitor cocktail (Roche), 1x PhosSTOP Phosphatase Inhibitor Cocktail (Roche) and 50µM MG132 ^44^. Protein concentration was quantified using and following the protocol of a BCA protein assay kit (Pierce). Protein concentrations were equalized by dilution with RIPA buffer and incubated for 10 minutes with loading buffer (5x sample loading buffer, Bio Rad) + β-ME) at 70°C before loading (30 µg total protein per sample) on pre-cast Mini-PROTEAN Stain Free TGX Gels (Bio Rad) and ran by SDS-PAGE. Gels were imaged before and after transferring to PVDF membranes (Bio Rad) using trans-blot turbo transfer system (Bio Rad), to verify successful and equal protein transfer. Blots were blocked for at least 1 hour in blocking solution at RT (5% milk in 1xTBS) before probing with primary antibody in blocking solution (α-HA-HRP, 1:2500 (Roche); α-PGB1, 1:500 (produced for this study using full length protein as antigen by GenScript); α-Actin, 1:2500 (Thermo Scientific) overnight at 4°C. Blots were rinsed 3 times with 1xTBS-T (0.1% Tween 20) for 10 minutes under gentle agitation before probing with secondary antibody (α-rabbit IgG-HRP for PGB1, 1:3000; α-mouse IgG-HRP for Actin, 1:2500) and/or SuperSignal™ West Femto chemiluminescence substrate (Fisher Scientific) and blot imaging using Image Lab software in a chemi-gel doc (Bio-rad) with custom accumulation sensitivity settings for optimal contrast between band detection and background signal. To visualize RAP2.3 (~45 kDa) and ACTIN (~42 kDa) protein levels on the same blot, membranes were stripped after taking final blot images using a mild stripping buffer (pH 2,2, 1.5% (w/v) glycine, 0.1% SDS and 1.0% Tween 20) for 15 minutes and rinsed 3x in 1xTBS-T before blocking and probing with the 2^nd^ primary antibody of interest.

### NO quantification

Intracellular NO levels were visualized using DAF-FM diacetate (7’-difluorofluorescein diacetate, Bio-Connect). Seedlings were incubated in the dark for 15 min under gentle agitation in 10mM Tris-HCl buffer (pH 7.4) containing 50μM DAF-FM DA and subsequently washed twice for 5 min 10mM Tris-HCl buffer (pH 7.4). Several roots of all treatments/genotypes were mounted in 10mM Tris-HCl buffer (pH 7.4) on the same microscope slide. Fluorescence was visualized using a Zeiss Observer Z1 LSM700 confocal microscope (oil immersion, 40x objective Plan-Neofluar N.A. 1.30) with excitation at 488 nm and emission at 490-555 nm. Roots incubated and mounted in 10mM Tris-HCl buffer (pH 7.4) without DAF-FM DA were used to set background values where no fluorescence was detected. Within experiments, laser power, pinhole, digital gain and detector offset were identical for all samples. Mean DAF-FM DA fluorescence pixel intensity in root tips was determined in similar areas of ~17000 μm^2^ between epidermis layers using ICY software (http://icy.bioimageanalysis.org/).

### Confocal Microscopy

Transgenic Arabidopsis seedlings of *35S:EIN3-GFP* and *35S:RAP2.12-GFP* and were fixed in 4% PFA (pH 6.9) right after treatments, kept under gentle agitation for 1h, were subsequently washed twice for 1 min in 1 x PBS and stored in ClearSee clearing solution (xylitol 10% (w/v), sodium deoxycholate 15% (w/v) and urea 25% (w/v) ^45^. Seedlings were transferred to 0.01% Calcofluor White (in ClearSee solution) 24 hours before imaging. Fluorescence was visualized using a Zeiss Observer Z1 LSM700 confocal microscope (oil immersion, 40x objective Plan-Neofluar N.A. 1.30) with excitation at 488nm and emission at 490-555nm for GFP and excitation at 405 nm and emission at 400-490 nm for Calcofluor White. Within experiments, laser power, pinhole, digital gain and detector offset were identical for all samples. Mean GFP fluorescence pixel intensity in root tips was determined in similar areas of ~17000 μm^2^ between epidermis layers using ICY software (http://icy.bioimageanalysis.org/).

### Nitrate reductase activity assay

The NR activity was assessed using a mix of 20 whole 10-day-old seedlings with 2 replicates per treatment. Snap frozen samples were ground and homogenized in extraction buffer (100mM HEPES (pH7.5), 2mM EDTA, 2mM di-thiothreitol, 1% PVPP). After centrifugation at 30.000g at 4C for 20 min, supernatants were collected and added to the reaction buffer (100mM HEPES (pH7.5), 100mM NaNO3, 10mM Cysteine, 2mM NADH and 2mM EDTA). The reaction was stopped by the addition of 500mM zinc acetate after incubation for 15min at 25°C. Total nitrite accumulation was determined following addition of 1% sulfanilamide in 1.5M HCl and 0.02% naphthylethylenediamine dihydrochloride (NNEDA) in 0.2M HCl by measuring the absorbance of the reaction mixture at 540 nm.

### Statistical analyses

No statistical methods were used to predetermine sample size. Samples were taken from independent biological replicates. In general, the sample size of experiments was maximized and dependent on technical, space and/or time limitations. For root tip survival assay, the maximum amount of seedlings used per biological replicate, generally 1 row of seedlings for in vitro agar plates, is mentioned in the appropriate figure legends. Data was plotted using Graphpad Prism software. The statistical tests were performed two-sided, using R software and the “LSmeans and “multmultcompView” packages. Surival data was analyzed with either a generalized linear modeling (GLM) approach or an ANOVA on arcsin transformation of the surviving fraction. A negative binomial error structure was used for the GLM. Arcsin transformation ensured a homogeneity and normal distribution of the variances, especially for data that did not have treatments with all living or all death responses. The remaining data were analyzed with either Students t-test, 1-way or 2-way ANOVAs. Here data were log transformed if necessary to adhere to ANOVA prerequisites. Multiple comparisons were corrected for with Tukey’s HSD.

## Acknowledgements

We thank the following individuals for providing seeds of these genotypes: Angelika Mustroph for Ler-0, Col-0 x Ler-0 WT crosses, *rap2.2*, *rap2.12* and *rap2.2 rap2.12-A* & *B*, Francesco Licausi for *35S:δ13-RAP2.12-GFP* and *35S:RAP2.12-GFP*, Shi Xiao for *35S:EIN3-GFP* and Frank Becker for *Arabidopsis lyrata* seeds. We acknowledge Sophie Berckhan, Ankie Ammerlaan, Rob Welschen, Tamara Le Thanh, Johanna Kociemba, Florian de Deugd and Joris te Riele for technical assistance. Finally, we thank Ronald Pierik for feedback on the manuscript and Kasper van Gelderen and Jesse Küpers for their input on confocal imaging. This work was supported by grants from the Netherlands Organization for Scientific Research (831.15.001 to S.H., 824.14.007 to L.A.C.J.V, S.M. and BB.00534.1 to R.S.) and the Biotechnology and Biological Sciences Research Council [BB/M007820/1 and BB/K000144/1] to M.J.H.

## Author information and contributions

Authors declare no competing interests. S.H, Z.L, H.v.V, J.V.C, H.Z, E.J.W.V, J.B.S, F.L.T, K.H.H, D.J.G, M.J.H, R.S. and L.A.C.J.V. designed experiments; S.H, Z.L, J.V.C, E.R, S.M, H.Z, N.v.D, F.B, G.W.B and E.J.W.V performed experiments; S.H, M.J.H, R.S. and L.A.C.J.V. wrote the manuscript.

## Data and materials availability

No restrictions are placed on materials and data availability. Biological materials such as mutant/transgenic lines can be requested from the corresponding authors. Details of all data and materials used in the analysis are available in the main text or the supplementary materials. Gene accession numbers of all the Arabidopsis genes/mutants used in this study are listed in the Method section and Supplementary Table 1 and 2.

## Supplementary Information

Supplementary files include:

Supplementary figures.

Supplementary Tables 1 and 2. List of accessions used with genotyping primers (Supplementary Table 1) and RT-qPCR primers (Supplementary Table 2).

## Supplementary Figures

**Supplementary Figure 1.**
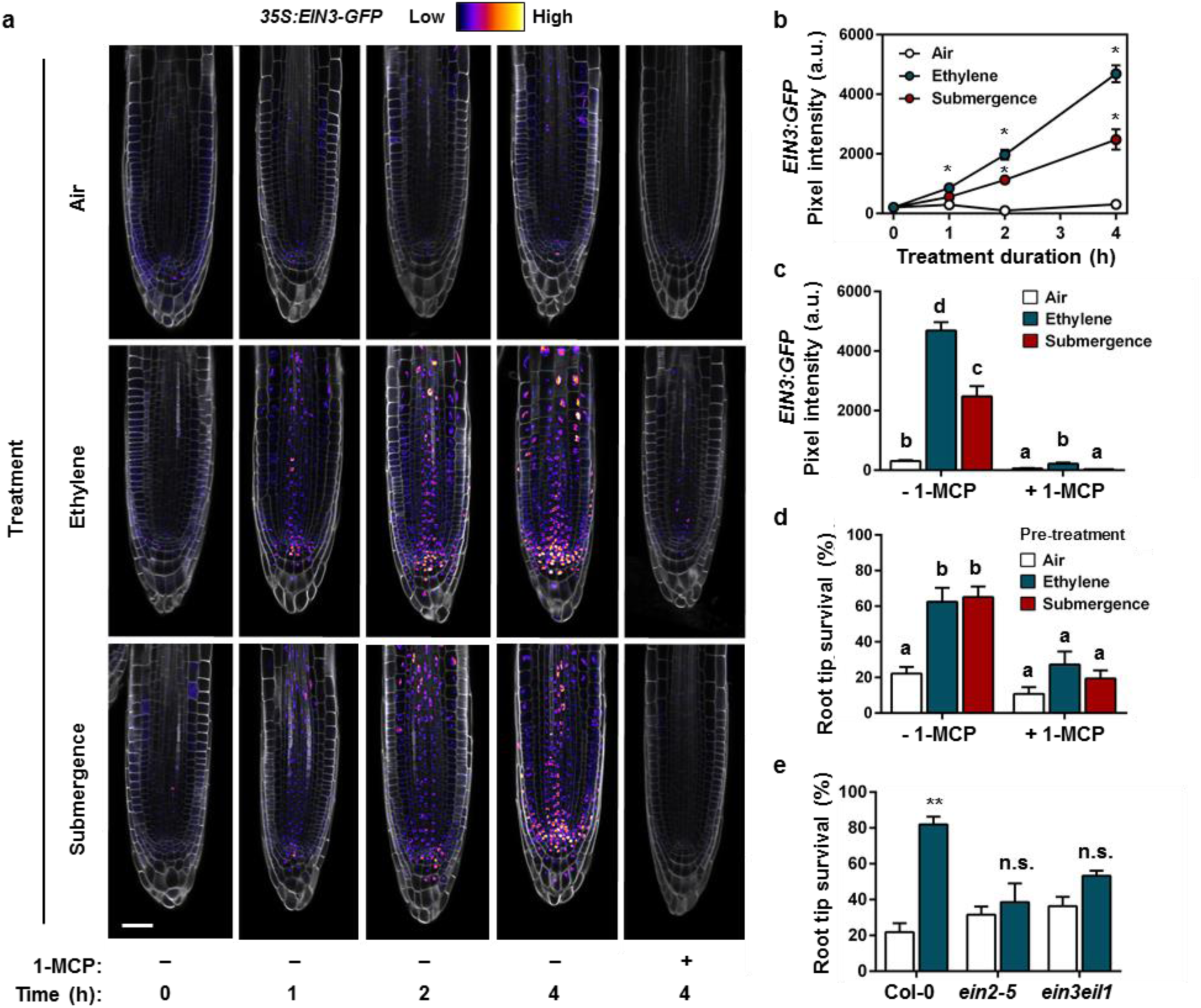
Ethylene signalling upon early submergence and its functional implications for subsequent hypoxia acclimation. **(a)** Representative confocal images of protein stability and localization of the ethylene master regulator EIN3, using the *35S:EIN3-GFP* (*ein3eil1* double mutant background) signal in Arabidopsis root tips. Seedling were treated for up to 4 hours with air, ~5μll^−1^ ethylene or submergence, either in combination with or without a pre-treatment of ethylene action inhibitor 1-MCP at the 4 hour time-point. Cell walls were visualized using Calcofluor White stain (scale bar= 50μm). **(b)** Quantification of *35S::EIN3-GFP* in root tips during 4 hours of treatment with air (white), ~5μll^−1^ ethylene (blue) or submergence (red). Mean GFP pixel intensity inside the root tips was quantified using ICY imaging software. Asterisks indicate a significant difference from the air mean per time-point (Error bars are SEM, p<0.05, ANOVA with planned comparisons, Tukey’s HSD correction for multiple comparisons, n=5-12 roots). **(c)** Quantification of *35S::EIN3-GFP* (*ein3eil1* double mutant background) in root tips after 4 hours of treatment with air (white), ~5μll^−1^ ethylene (blue) or submergence (red), either in combination with or without a pre-treatment of ethylene action inhibitor 1-MCP. Mean GFP pixel intensity inside the root tips was quantified using ICY imaging software. Samples without 1-MCP are the same as in Supplementary Figure. 1b at t=4h. Statistically similar groups are indicated using the same letter (Error bars are SEM, p<0.05, 2-way ANOVA, Tukey’s HSD, n=5-11 roots). **(d)** Seedling root tip survival of Col-0 after 4 hours of pre-treatment with air (white), ~5μll^−1^ ethylene (blue) or submergence (red), either in combination with or without a pre-treatment of ethylene action inhibitor 1-MCP, followed by 4 hours of hypoxia and 3 days of recovery. Values are relative to control (normoxia) plants. Statistically similar groups are indicated using the same letter (Error bars are SEM, p<0.05, Generalized linear model with negative binomial error structure, n=8 rows of ~23 seedlings). **(e)** Seedling root tip survival of Col-0 and two ethylene signaling pathway loss-of-function mutants after 4 hours of pre-treatment with air (white) or ~5μll^−1^ ethylene (blue) followed by 4 hours of hypoxia and 3 days of recovery. Values are relative to control (normoxia) plants. Asterisks indicate significant differences between air and ethylene (Error bars are SEM, p<0.01, Generalized linear model with negative binomial error structure, n=4-6 rows of ~46 seedlings). Experiments were replicated at least 2 times.

**Supplementary Figure 2.**
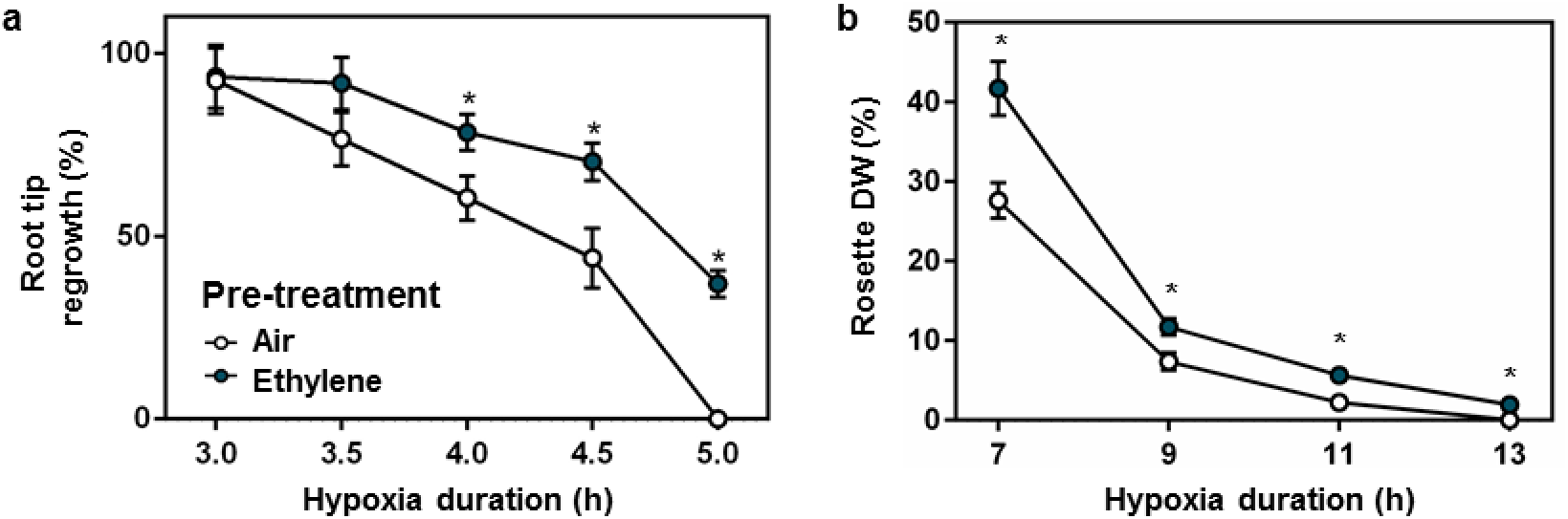
Ethylene pre-treatment improves performance of recovering tissues of survived plants after subsequent hypoxia. **(a)** Seedling root tip regrowth capacity of surviving roots after 4 hours of pre-treatment with air (white) or ~5μll^−1^ ethylene (blue) followed by hypoxia and 3 days of recovery. Values are relative to control (normoxia) plants. Asterisks indicate significant differences between air and ethylene at given time point (Error bars are SEM, *p<0.05, Student’s *t* test, n=4-8 lines of 23 seedlings for survival, n= 5-35 surviving roots for regrowth). **(b)** Rosette dry weight (DW) of adult Col-0 plants after 4 hours of pre-treatment with air (white) or ~5μll^−1^ ethylene (blue) followed by hypoxia and 7 days of recovery. DW was measured only from surviving plants. Values are relative to control (normoxia) plants. Asterisks indicate significant differences between air and ethylene at given time point (Error bars are SEM, *p<0.05, Student’s *t* test, n=30 plants). Experiments were replicated at least 3 times.

**Supplementary Figure 3.**
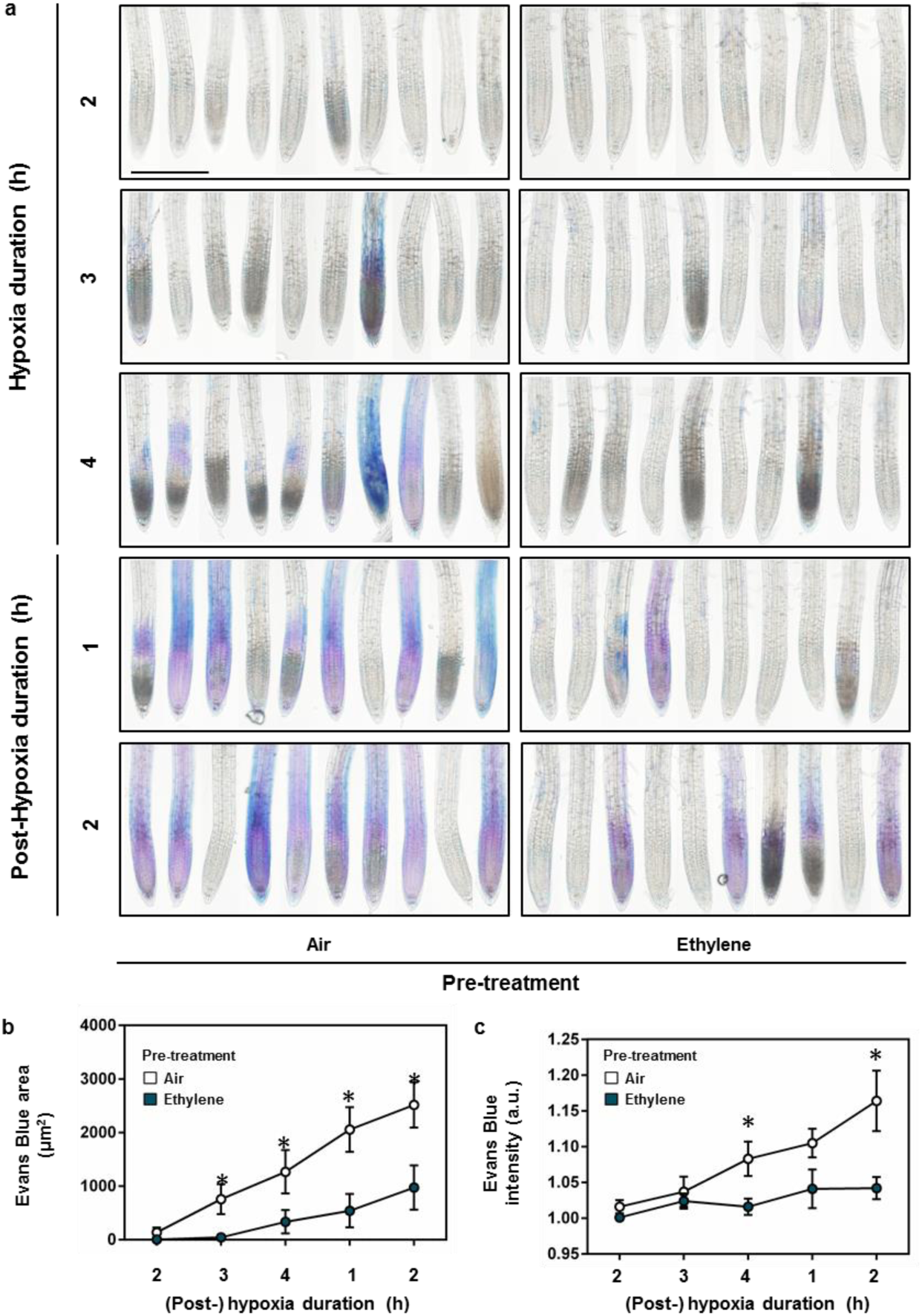
Ethylene pre-treated seedlings show reduced cell damage in root tips during subsequent hypoxia and recovery treatments. **(a)** Representative light microscopy images of Evans blue staining for impaired cell membrane integrity in seedling root tips after 4 hours of pre-treatment with air or ~5μll^−1^ ethylene followed by 2-4h hypoxia (scale bar = 2mm). **(b, c)** Quantification of the area (b) and pixel intensity (c) of Evans blue staining in seedling root tips after 4 hours of pre-treatment with air (white) or ~5μll^−1^ ethylene (blue) followed by 2-4h hypoxia and 1-2h of recovery. Asterisks indicate significant differences between air and ethylene at given time point (Error bars are SEM, *p<0.05, Student’s t test, n=10 root tips). Experiments were replicated at least 2 times.

**Supplementary Figure 4.**
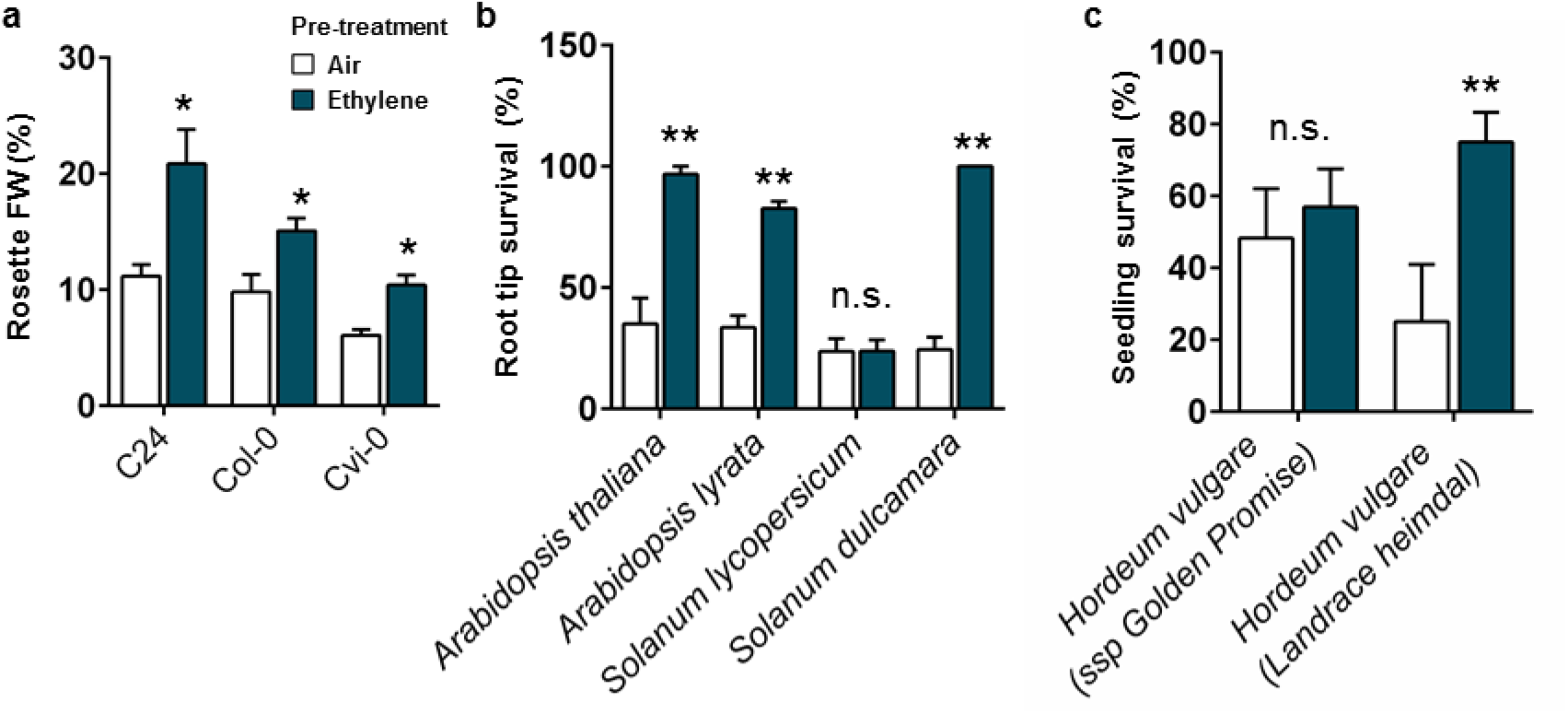
Ethylene-induced hypoxia tolerance is conserved within Arabidopsis accessions and shows variation between other plant species. **(a)** Relative rosette fresh weight (FW) of adult Arabidopsis accessions C24, Col-0 and Cvi-0 plants after 4 hours of pre-treatment with air (white) or ~5μll^−1^ ethylene (blue) followed by 9 hours of hypoxia and 7 days of recovery. FW was measured only from survived plants (Error bars are SEM, *p<0.05, Student’s *t* test, n=10 plants). **(b)** Root tip survival of 4 different plants species after 4 hours of pre-treatment with air (white) or ~5μll^−1^ ethylene (blue) followed by 4 hours of hypoxia and 3 days of recovery Asterisks indicate significant differences between air and ethylene (Error bars are SEM, **p<0.01, Generalized linear model with negative binomial error structure, n=4-6 lines consisting of 10-46 seedlings depending on species). **(c)** Plant survival of 2 different varieties of Barley (*Hordeum vulgare*) seedlings after 4 hours of pre-treatment with air (white) or ~5μll^−1^ ethylene (blue) followed by 20 hours of hypoxia and 3 days of recovery. Asterisks indicate significant differences between air and ethylene (Error bars are SEM, *p<0.05, Generalized linear model with negative binomial error structure, n=4-6 replicates consisting of 3 seedlings). Experiments were replicated at least 2 times, except for a, which was only performed once.

**Supplementary Figure 5.**
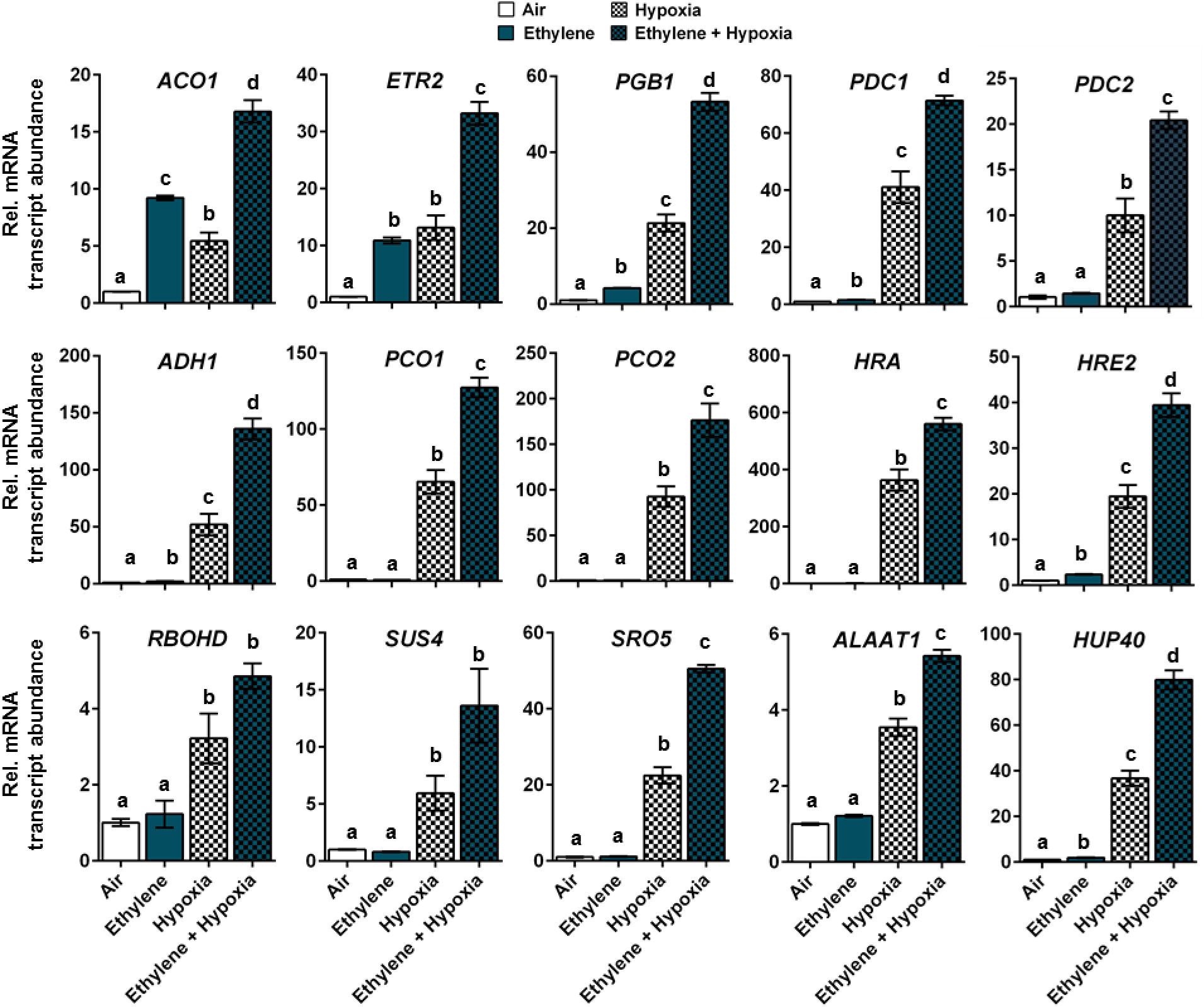
Ethylene pre-treatment bolsters hypoxia adaptive gene transcripts upon hypoxia. Relative mRNA transcript abundance of 15 hypoxia adaptive genes in seedling root tips of Col-0 after 4 hours of pre-treatment with air (white) or ~5μll^−1^ ethylene (blue), followed by (4h) hypoxia (blocks). Values are relative to Col-0 air treated samples. Different letters indicate significant differences (Error bars are SEM, p<0.05, 1-way ANOVA, Tukey’s HSD, n=3-4 replicates of ~400 root tips). Experiments were replicated at least 2 times.

**Supplementary Figure 6.**
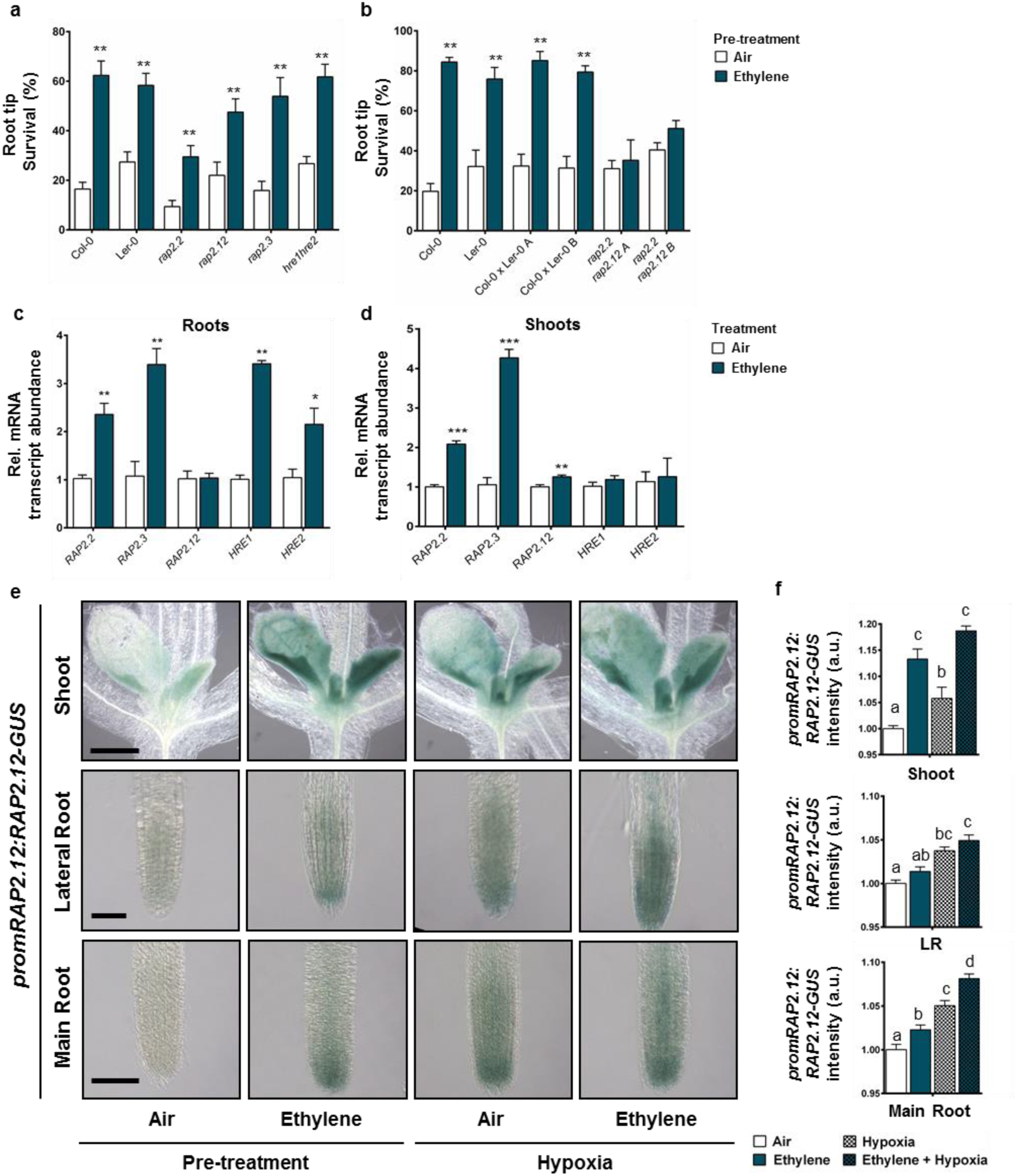
The involvement and regulation of ERFVIIs for ethylene-induced hypoxia tolerance. **(a, b)** Seedling root tip survival of Col-0, Ler-0, ERFVII mutants *rap2.2* (Ler-0 background), *rap2.12*, *rap2.3* and *hre1hre2* (Col-0 background) in a, and Col-0, Ler-0, 2 Col-0 x Ler-0 WT crosses and ERFVII double mutants *rap2.2rap2.12* (2 independent lines in Col-0 x Ler-0 background) in b, after 4 hours of pre-treatment with air (white) or ~5μll^−1^ ethylene (blue) followed by 4 hours of hypoxia and 3 days of recovery. Values are relative to control (normoxia) plants. Asterisks indicate significant differences between air and ethylene (Error bars are SEM, **p<0,01, Generalized linear model with negative binomial error structure, n=4-21 rows consisting of ~23 seedlings for a, n=8 rows consisting of ~23 seedlings for b). **(c, d)** Relative mRNA transcript abundance of all 5 ERFVIIs in root tips of Col-0 seedlings (c) and adult rosettes (d) after 4 hours of treatment with air (white) or ~5μll^−1^ ethylene (blue). Asterisks indicate significant differences between air and ethylene (Error bars are SEM,*p<0.05, **p<0.01, ***p<0.001, Generalized linear model with negative binomial error structure, n=3-4 replicates containing ~400 root tips for c, n=5 replicates of 2 rosettes for d). **(e, f)** Representative DIC microscopy images (e) and quantification (f) of *promRAP2.12::RAP2.12-GUS* in seedling shoots, lateral roots and main root tips after 4 hours of treatment with air (white) or ~5μll^−1^ ethylene (blue) or subsequent (4h) hypoxia (block pattern). Scale bars; shoot = 180μm, lateral root = 60μm, main root = 100μm. Values are relative to air treated samples. Statistically similar groups are indicated using the same letter per tissue (Error bars are SEM, p<0.05, 1-way ANOVA, Tukey’s HSD, n=5-20 replicates). Experiments were replicated at least 2 times.

**Supplementary Figure 7.**
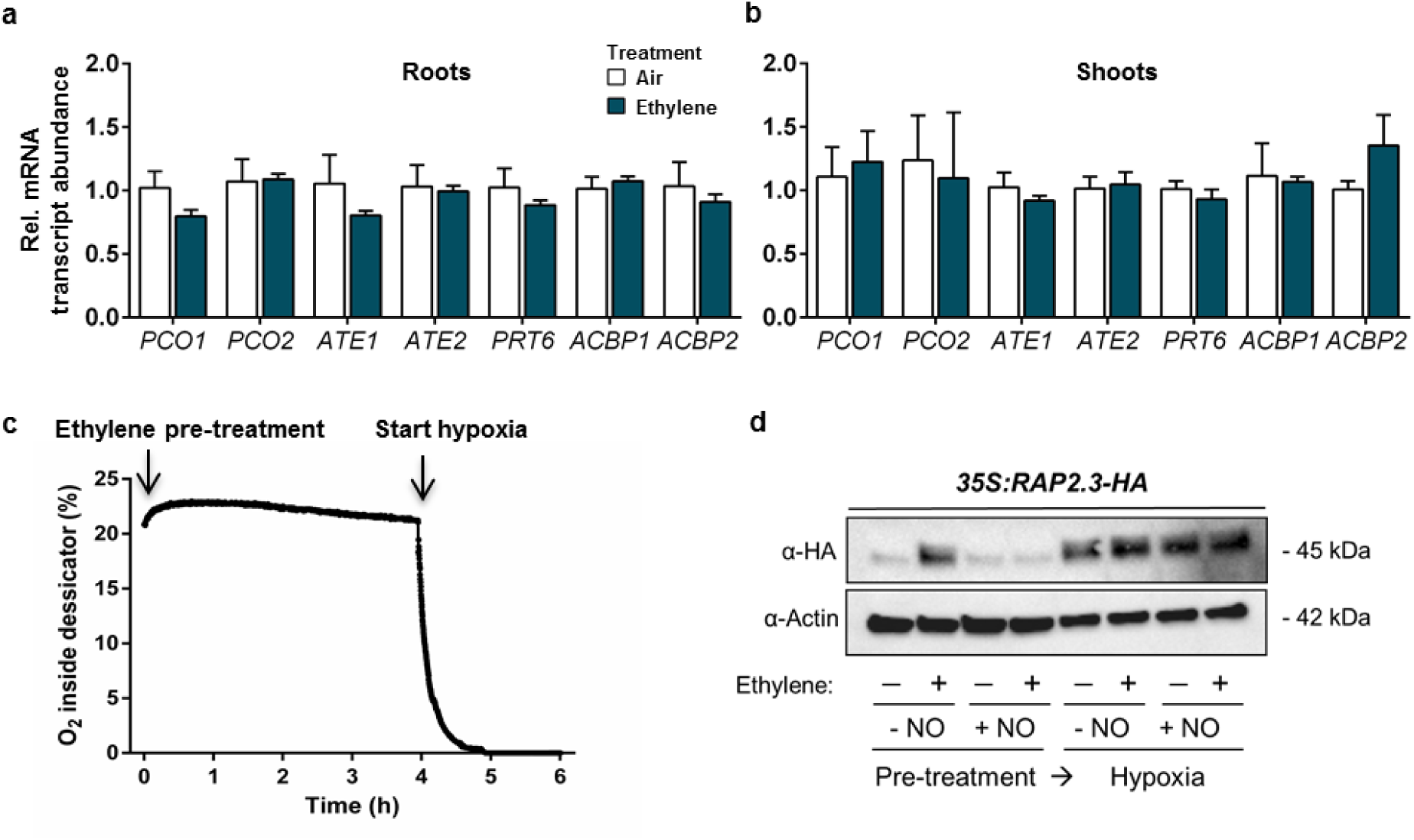
The effects of ethylene on processes that regulate ERFVII stability. **(a, b)** Relative mRNA transcript abundance of genes coding for enzymes involved in the PRT6 N-degron pathway or RAP2.12-sequestering proteins *ACBP1* and *ACBP2* in root tips of Col-0 seedlings (a) and adult rosettes (b) after 4 hours of treatment with air (white) or ~5μll^−1^ ethylene (blue). Values are relative to Col-0 air treated samples. No significant differences were found between air and ethylene (Error bars are SEM, Student’s t test, n=3-4 replicates containing ~400 root tips for a, n=5 replicates of 2 rosettes for b). **(c)** Levels of molecular oxygen measured over time at the outflow of the desiccators during the ethylene pre-treatment and subsequent hypoxia treatments in this study. Oxygen levels generally reached <0.00% between 40 and 50 minutes of flushing the desiccators with humidified 99.996% N_2_ at a rate of 2l min^−1^. **(d)** RAP2.3 protein levels in 35S::MC-RAP2.3-HA seedlings (Col-0 background) after air and ethylene pre-treatments (4h), combined with or without an additional NO pulse and subsequent hypoxia (4h). Experiments were replicated at least 2 times, except for d, in which the hypoxia treatment after NO manipulation was only performed once.

**Supplementary Figure 8.**
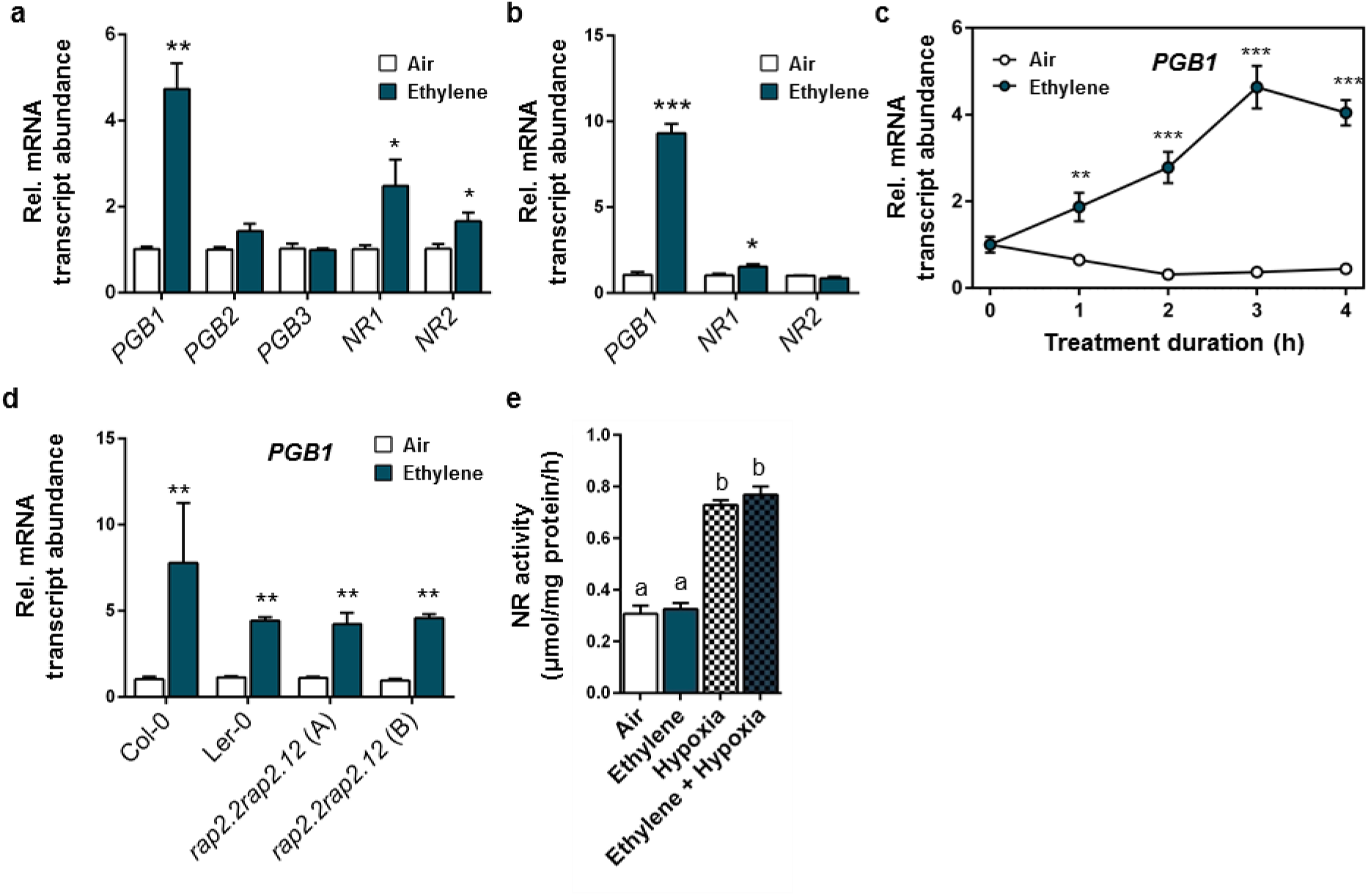
The effects of ethylene on *PHYTOGLOBIN* transcript abundance and *NITRATE REDUCTASE* transcript abundance and activity. **(a)** Relative mRNA transcript abundance of genes involved in NO metabolism in seedling root tips of Col-0 after 4 hours of treatment with air (white) or ~5μll^−1^ ethylene (blue). Values are relative to Col-0 air treated samples. Asterisks indicate significant differences between air and ethylene (Error bars are SEM, *p<0.05, **p<0.01, Student’s *t* test, n=3-4 biological replicates of ~00 root tips). **(b)** Relative mRNA transcript abundance of genes coding for enzymes involved in NO metabolism in rosettes of Col-0 plants after 4 hours of treatment with air (white) or ~5μll^−1^ ethylene (blue). Values are relative to Col-0 air treated samples. Asterisks indicate significant differences between air and ethylene (Error bars are SEM, ***p<0,001, *p<0,05 Student’s *t* test, n=5 biological replicates of 2 rosettes). **(c)** Relative *PGB1* mRNA transcript abundance in rosettes of Col-0 plants during 4 hours of treatment with air (white) or ~5μll^−1^ ethylene (blue). Asterisks indicate significant differences between air and ethylene (Error bars are SEM, ***p<0,001, ANOVA with planned comparisons, Tukey’s HSD correction for multiple comparisons, n=5 biological replicates of 2 rosettes). **(d)** Relative *PGB1* mRNA transcript abundance in seedlings of Arabidopsis Col-0 and Ler-0 WT, and 2 double *rap2.2rap2.12* mutants (Col-0 x Ler-0 background) after 4 hours of treatment with air (white) or ~5μll^−1^ ethylene (blue). Values are relative to Col-0 air treated samples. Asterisks indicate significant differences between air and ethylene (Error bars are SEM, **p<0.01, ANOVA with planned comparisons, Tukey’s HSD correction for multiple comparisons, n=2 biological replicates of ~400 root tips). **(e)** Nitrate reductase activity in whole Col-0 WT seedlings after 4 hours of pre-treatment with air (white) or ~5μll^−1^ ethylene (blue), followed by (4h) hypoxia (blocks). Statistically similar groups are indicated using the same letter (Error bars are SEM, p<0.05, 1-way ANOVA, Tukey’s HSD, n=2 biological replicates of ~200 seedlings). Experiments were replicated at least 2 times, except for e, which was only performed once.

**Supplementary Figure 9.**
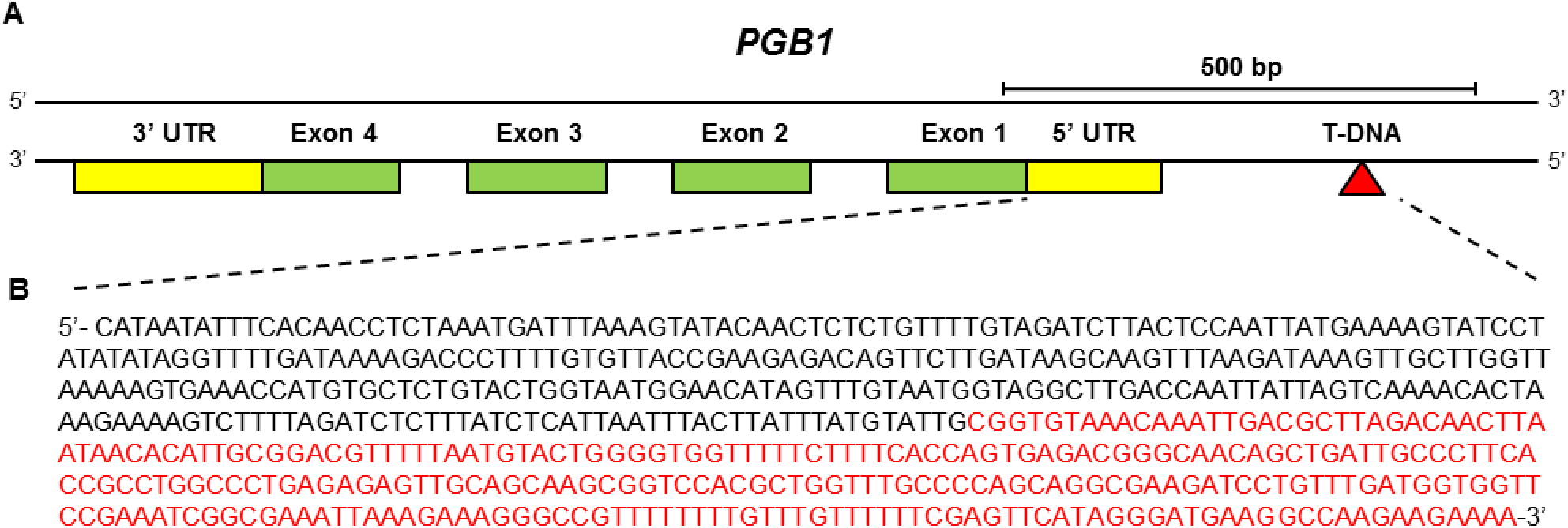
Identification of *pgb1-1* mutant line *SK_058388*. **(a)** Schematic map of genomic *PGB1* gene region including the 4 *PGB1* exons (green) and the location of the T-DNA insertion (red triangle) of *pgb1-1* line *SK_058388*. **(b)** Partial DNA sequencing reaction of *pgb1-1* aligned with genomic *PGB1* gene region. The aligned native *PGB1* sequence (black) ends and the T-DNA sequence (red) starts exactly 300bp upstream of the *PGB1* start codon.

**Supplementary Figure 10.**
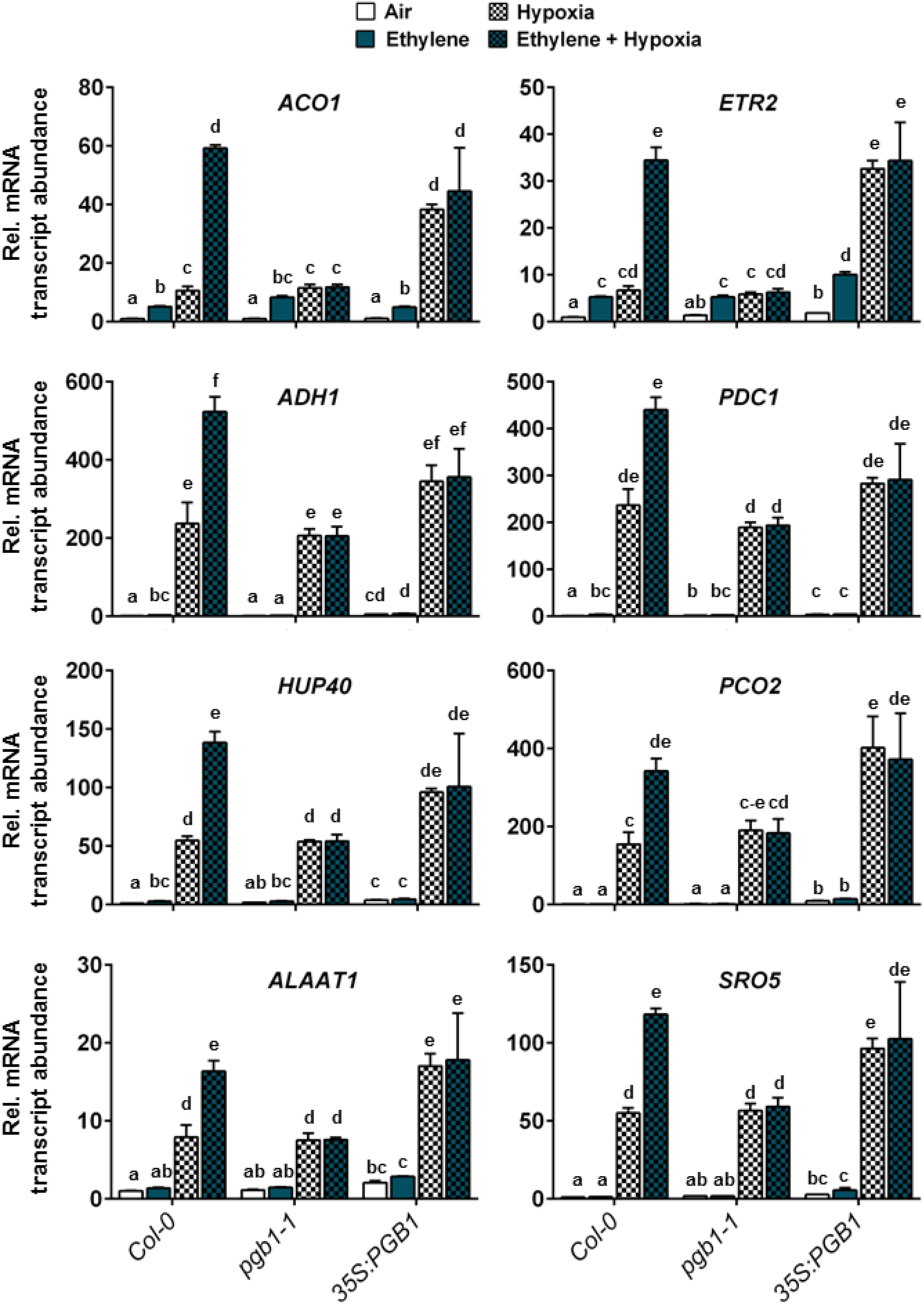
Hypoxia adaptive gene expression in *PGB1* knock-down and over-expression lines. Relative mRNA transcript abundance of 8 hypoxia adaptive genes in seedling root tips of Col-0*, pgb1-1* and *35S::PGB1* after 4 hours of pre-treatment with air (white) or ~5μll^−1^ ethylene (blue), followed by (4h) hypoxia (blocks). Values are relative to Col-0 air treated samples. Different letters indicate significant differences (Error bars are SEM, p<0.05, 2-way ANOVA, Tukey’s HSD, n=3 replicates of ~200 root tips).

**Table S1.**
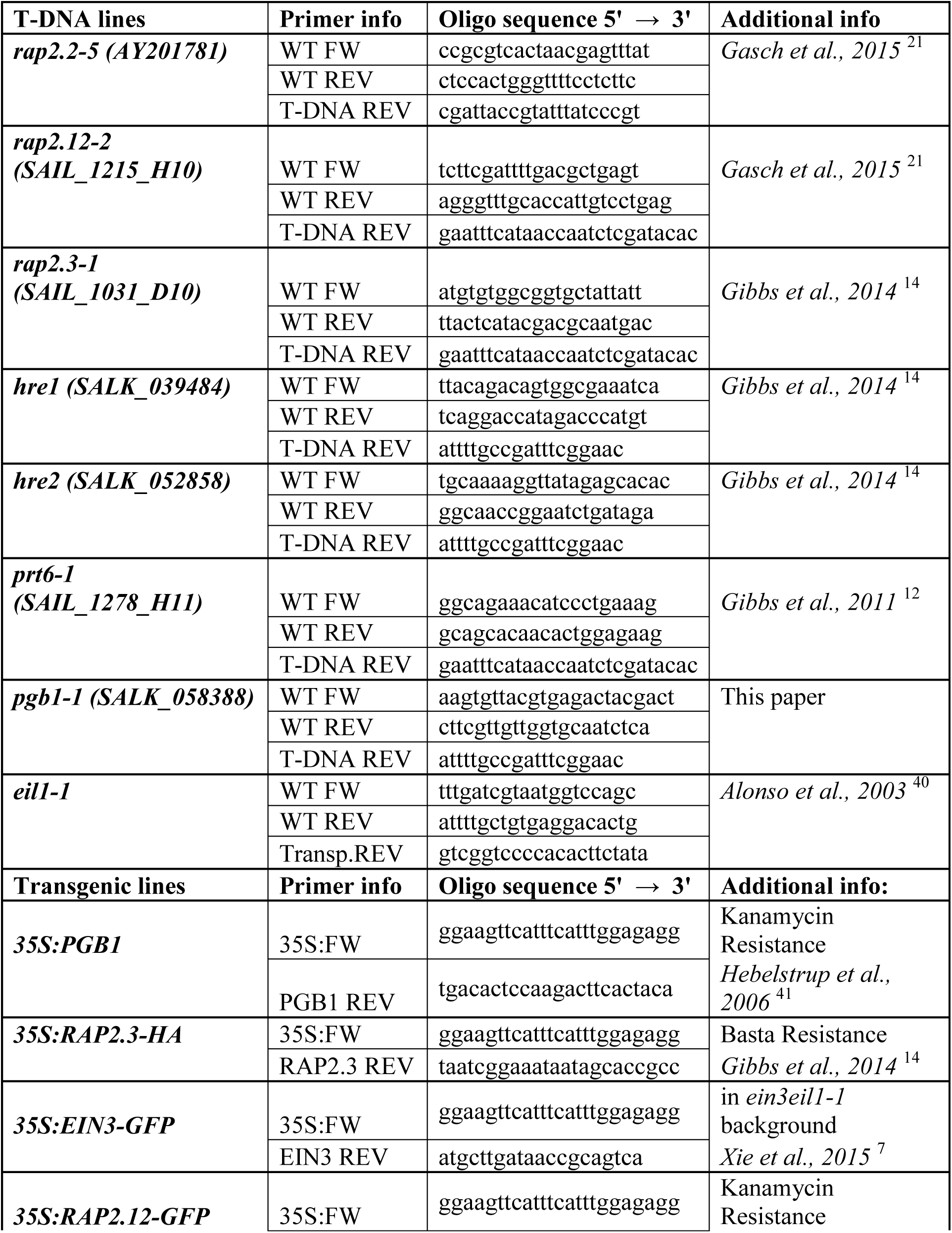
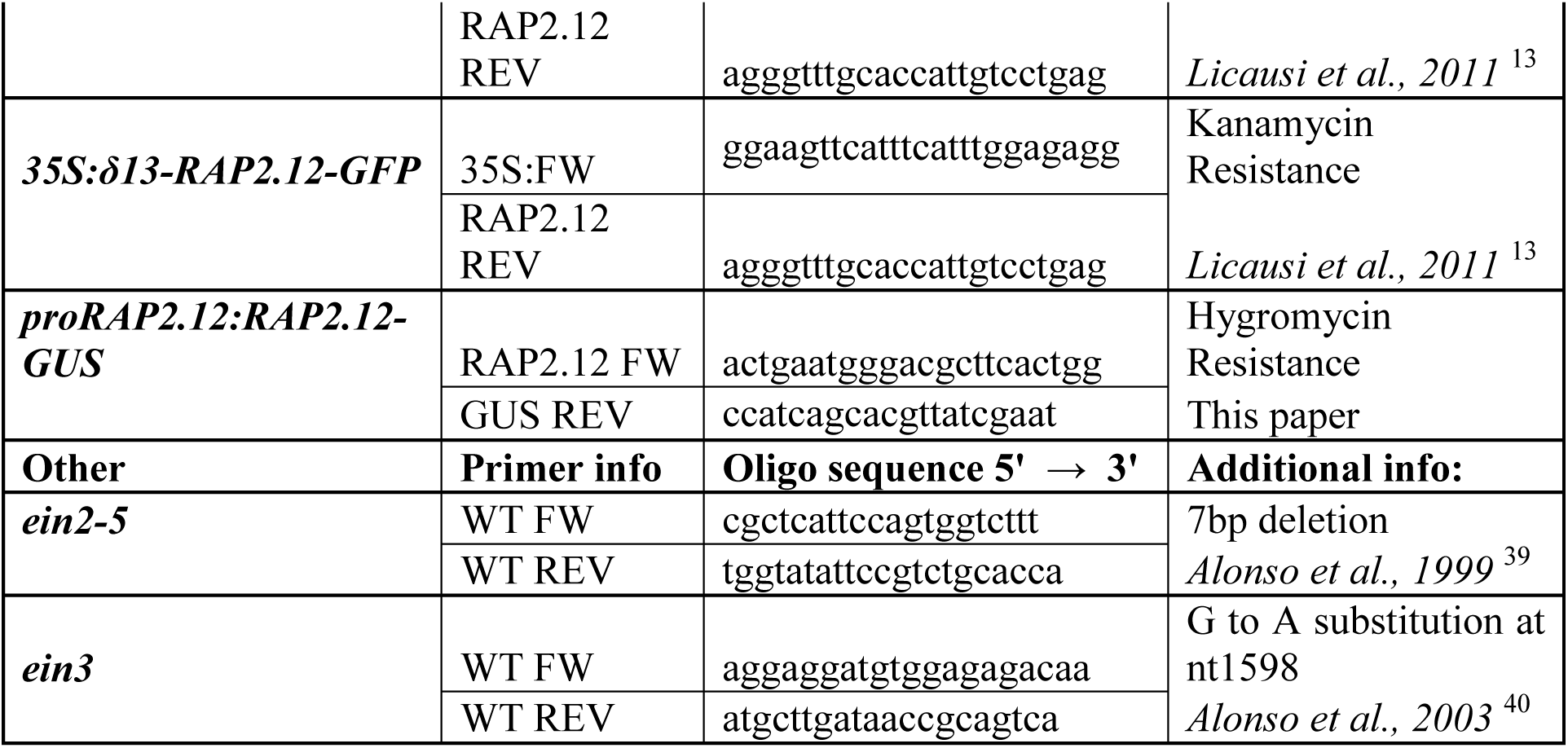
List of genotyping primers used in this study.

**Table S2.**
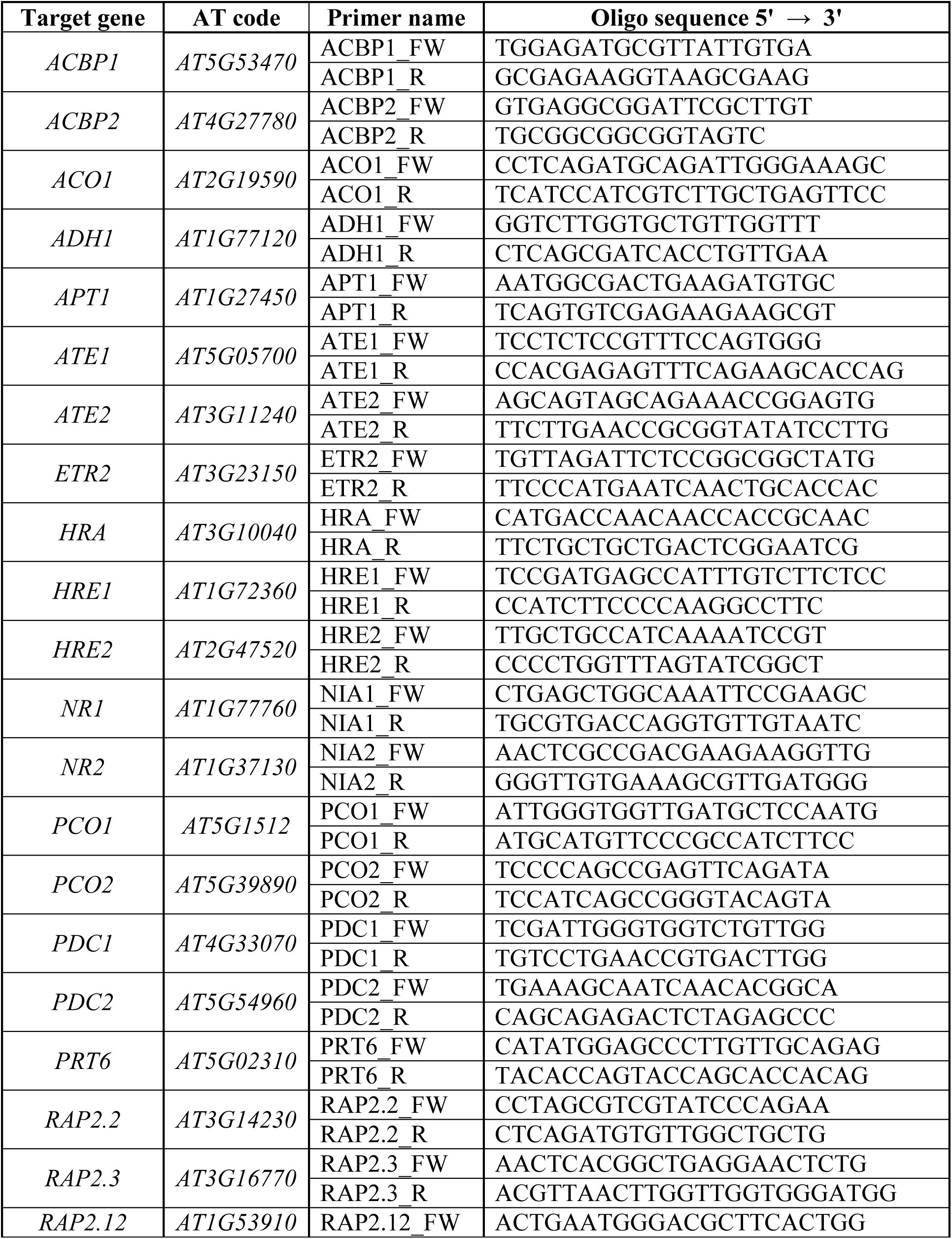
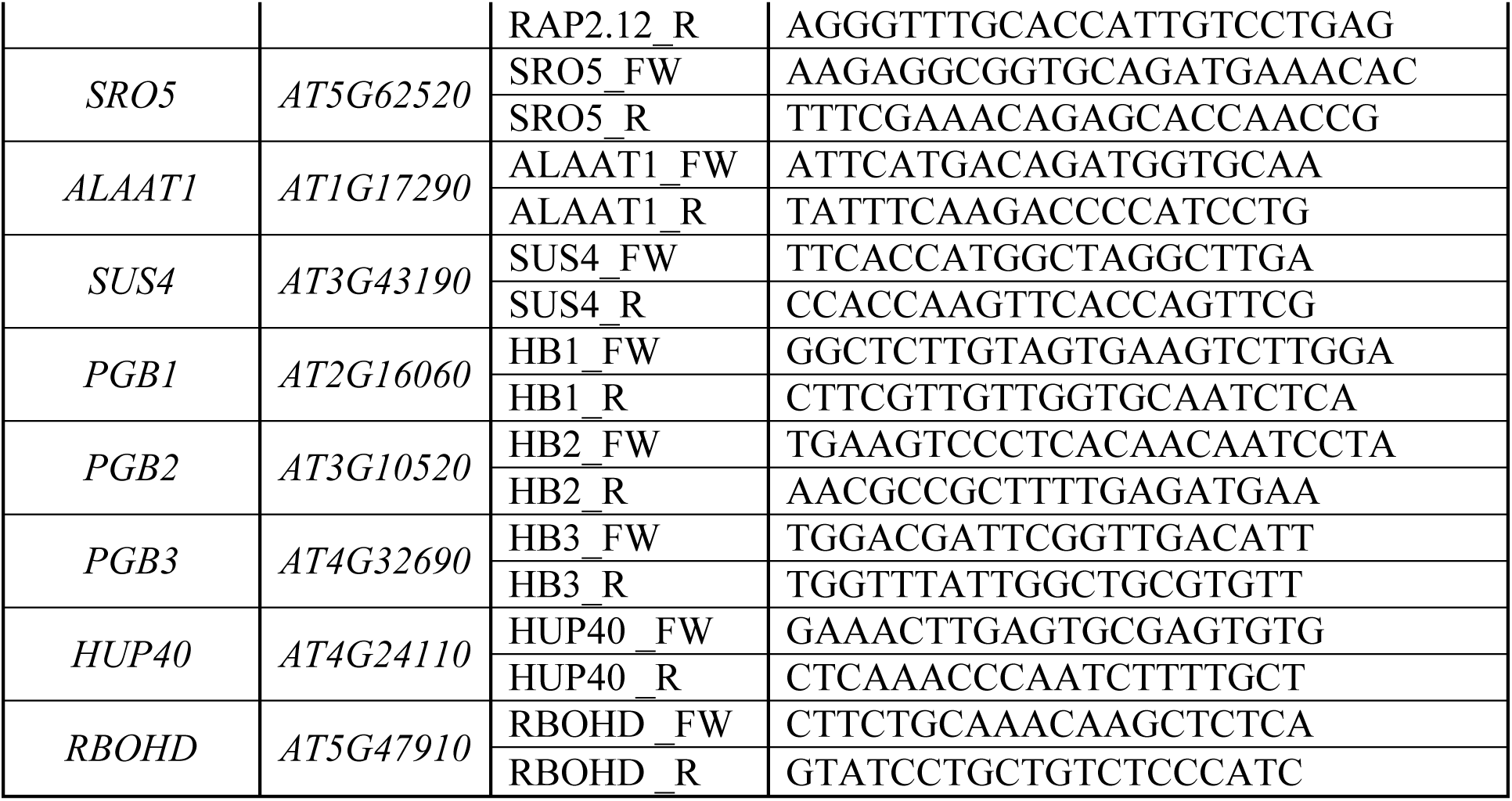
List of RT-qPCR primers used in this study.

## References

1. Hirabayashi, Y. et al. Global flood risk under climate change. Nat. Clim. Chang. 3, 816–821 (2013).

2. Voesenek, L. A. C. J. & Bailey-Serres, J. Flood adaptive traits and processes: an overview. New Phytol. 206, 57–73 (2015).

3. Shiono, K., Takahashi, H., Colmer, T. D. & Nakazono, M. Role of ethylene in acclimations to promote oxygen transport in roots of plants in waterlogged soils. Plant Sci. 175, 52–58 (2008).

4. Sasidharan, R. et al. Signal Dynamics and Interactions during Flooding Stress. Plant Physiol. 176, 1106–1117 (2018).

5. Voesenek, L. A. C. J. & Sasidharan, R. Ethylene - and oxygen signalling - drive plant survival during flooding. Plant Biol. 15, 426–435 (2013).

6. Chen, Y.-F., Etheridge, N. & Schaller, G. E. Ethylene Signal Transduction. Ann. Bot. 95, 901–915 (2005).

7. Xie, L. J. et al. Unsaturation of Very-Long-Chain Ceramides Protects Plant from Hypoxia-Induced Damages by Modulating Ethylene Signaling in Arabidopsis. PLoS Genet. 11, 1–33 (2015).

8. Chang, K. N. et al. Temporal transcriptional response to ethylene gas drives growth hormone cross-regulation in Arabidopsis. Elife 2, e00675 (2013).

9. An, F. et al. Ethylene-Induced Stabilization of ETHYLENE INSENSITIVE3 and EIN3-LIKE1 Is Mediated by Proteasomal Degradation of EIN3 Binding F-Box 1 and 2 That Requires EIN2 in Arabidopsis. Plant Cell Online 22, 2384–2401 (2010).

10. Veen, H. van et al. Two Rumex Species from Contrasting Hydrological Niches Regulate Flooding Tolerance through Distinct Mechanisms. Plant Cell 25, 4691–4707 (2013).

11. Mustroph, A. et al. Cross-Kingdom Comparison of Transcriptomic Adjustments to Low-Oxygen Stress Highlights Conserved and Plant-Specific Responses. Plant Physiol. 152, 1484–1500 (2010).

12. Gibbs, D. J. et al. Homeostatic response to hypoxia is regulated by the N-end rule pathway in plants. Nature 479, 415–418 (2011).

13. Licausi, F. et al. Oxygen sensing in plants is mediated by an N-end rule pathway for protein destabilization. Nature 479, 419–422 (2011).

14. Gibbs, D. J. et al. Nitric Oxide Sensing in Plants Is Mediated by Proteolytic Control of Group VII ERF Transcription Factors. Mol. Cell 53, 369–379 (2014).

15. White, M. D. et al. Plant cysteine oxidases are dioxygenases that directly enable arginyl transferase-catalysed arginylation of N-end rule targets. Nat. Commun. 8, 14690 (2017).

16. Tasaki, T., Sriram, S. M., Park, K. S. & Kwon, Y. T. The N-End Rule Pathway. Annu. Rev. Biochem. 81, 261–289 (2012).

17. Gibbs, D. J. et al. Group VII Ethylene Response Factors Coordinate Oxygen and Nitric Oxide Signal Transduction and Stress Responses in Plants. Plant Physiol. 169, 23–31 (2015).

18. Licausi, F. et al. Oxygen sensing in plants is mediated by an N-end rule pathway for protein destabilization. Nature 479, 419–422 (2011).

19. Vicente, J. et al. The Cys-Arg/N-End Rule Pathway Is a General Sensor of Abiotic Stress in Flowering Plants. Curr. Biol. 27, 3183–3190.e4 (2017).

20. Hinz, M. et al. Arabidopsis RAP2.2: An Ethylene Response Transcription Factor That Is Important for Hypoxia Survival. Plant Physiol. 153, 757–772 (2010).

21. Gasch, P. et al. Redundant ERF-VII Transcription Factors Bind to an Evolutionarily Conserved cis-Motif to Regulate Hypoxia-Responsive Gene Expression in Arabidopsis. Plant Cell 28, 160–180 (2016).

22. Bui, L. T., Giuntoli, B., Kosmacz, M., Parlanti, S. & Licausi, F. Constitutively expressed ERF-VII transcription factors redundantly activate the core anaerobic response in Arabidopsis thaliana. Plant Sci. 236, 37–43 (2015).

23. Licausi, F. et al. HRE1 and HRE2, two hypoxia-inducible ethylene response factors, affect anaerobic responses in Arabidopsis thaliana. Plant J. 62, 302–315 (2010).

24. Mendiondo, G. M. et al. Enhanced waterlogging tolerance in barley by manipulation of expression of the N-end rule pathway E3 ligase PROTEOLYSIS6. Plant Biotechnol. J. 14, 40–50 (2016).

25. Weits, D. A. et al. An apical hypoxic niche sets the pace of shoot meristem activity. Nature 569, 714–717 (2019).

26. Shukla, V. et al. Endogenous Hypoxia in Lateral Root Primordia Controls Root Architecture by Antagonizing Auxin Signaling in Arabidopsis. Mol. Plant 12, 538–551 (2019).

27. Gibbs, D. J. et al. Oxygen-dependent proteolysis regulates the stability of angiosperm polycomb repressive complex 2 subunit VERNALIZATION 2. Nat. Commun. 9, 5438 (2018).

28. Planchet, E. & Kaiser, W. M. Nitric oxide (NO) detection by DAF fluorescence and chemiluminescence: A comparison using abiotic and biotic NO sources. J. Exp. Bot. 57, 3043–3055 (2006).

29. Gupta, K. J., Hebelstrup, K. H., Mur, L. A. J. & Igamberdiev, A. U. Plant hemoglobins: Important players at the crossroads between oxygen and nitric oxide. FEBS Lett. 585, 3843–3849 (2011).

30. Chamizo-Ampudia, A., Sanz-Luque, E., Llamas, A., Galvan, A. & Fernandez, E. Nitrate Reductase Regulates Plant Nitric Oxide Homeostasis. Trends Plant Sci. 22, 163–174 (2017).

31. Hebelstrup, K. H. et al. Haemoglobin modulates NO emission and hyponasty under hypoxia-related stress in Arabidopsis thaliana. J. Exp. Bot. 63, 5581–5591 (2012).

32. Loreti, E., Valeri, M. C., Novi, G. & Perata, P. Gene Regulation and Survival under Hypoxia Requires Starch Availability and Metabolism. Plant Physiol. 176, 1286–1298 (2018).

33. Schmidt, R. R. et al. Low-oxygen response is triggered by an ATP-dependent shift in oleoyl-CoA in Arabidopsis. Proc. Natl. Acad. Sci. U. S. A. 115, E12101–E12110 (2018).

34. Igamberdiev, A. U. & Hill, R. D. Elevation of cytosolic Ca2+ in response to energy deficiency in plants: the general mechanism of adaptation to low oxygen stress. Biochem. J. 475, 1411–1425 (2018).

35. Holdsworth, M. J. First hints of new sensors. Nat. Plants 1–2 (2017). doi:10.1038/s41477-017-0031-7

36. Mira, M. M., Hill, R. D. & Stasolla, C. Phytoglobins Improve Hypoxic Root Growth by Alleviating Apical Meristem Cell Death. Plant Physiol. 172, 2044–2056 (2016).

37. Rivera-Contreras, I. K. et al. Transcriptomic analysis of submergence-tolerant and sensitive Brachypodium distachyon ecotypes reveals oxidative stress as a major tolerance factor. Sci. Rep. 6, 1–15 (2016).

38. Armstrong, W., Beckett, P. M., Colmer, T. D., Setter, T. L. & Greenway, H. Tolerance of roots to low oxygen: ‘anoxic’ cores, the phytoglobin-nitric oxide cycle, and energy or oxygen sensing. J. Plant Physiol. (2019). doi:10.1016/J.JPLPH.2019.04.010

39. Alonso, J. M., Hirayama, T., Roman, G., Nourizadeh, S. & Ecker, J. R. EIN2, a bifunctional transducer of ethylene and stress responses in Arabidopsis. Science (80-.). (1999). doi:10.1126/science.284.5423.2148

40. Alonso, J. M. et al. Five components of the ethylene-response pathway identified in a screen for weak ethylene-insensitive mutants in Arabidopsis. Proc. Natl. Acad. Sci. U. S. A. 100, 2992–7 (2003).

41. Hebelstrup, K. H., Hunt, P., Dennis, E., Jensen, S. B. & Jensen, E. Ø. Hemoglobin is essential for normal growth of Arabidopsis organs. Physiol. Plant. 127, 157–166 (2006).

42. Nakagawa, T. et al. Improved Gateway Binary Vectors: High-Performance Vectors for Creation of Fusion Constructs in Transgenic Analysis of Plants. Biosci. Biotechnol. Biochem. 71, 2095–2100 (2007).

43. Livak, K. J. & Schmittgen, T. D. Analysis of Relative Gene Expression Data Using Real-Time Quantitative PCR and the 2−ΔΔCT Method. Methods 25, 402–408 (2001).

44. Zhang, H. et al. N-terminomics reveals control of Arabidopsis seed storage proteins and proteases by the Arg/N-end rule pathway. New Phytol. 218, 1106–1126 (2018).

45. Ursache, R., Andersen, T. G., Marhavý, P. & Geldner, N. A protocol for combining fluorescent proteins with histological stains for diverse cell wall components. Plant J. 93, 399–412 (2018).

